# The amblyopic acuity deficit: impact on the identification of letters distorted by spatial scrambling algorithms

**DOI:** 10.1101/2025.07.02.662570

**Authors:** Raffles X Zhu, Robert F Hess, Alex S Baldwin

## Abstract

The letter acuity impairment in the amblyopic eye often exceeds predictions made from the cut-off spatial frequency for grating detection. Spatial scrambling in the amblyopic eye’s projections to the visual cortex has been proposed to bear some responsibility for this additional deficit. Using a novel stimulus algorithm that creates spatially scrambled bandpass letters, we generated stimuli simulating either: i) “cortical scrambling” at the output of oriented model “simple cells”, or ii) “subcortical scrambling” of isotropic subunits that combine to form these simple cells. We also investigated a more conventional “noise masking” with bandpass noise. We performed two bandpass letter identification experiments, equating the stimuli shown to each eye by normalising either: i) their contrast, presenting them at four times their contrast detection threshold; or ii) their spatial scale, presenting them at twice the participant’s acuity threshold for each eye. At the group level, we found that the amblyopic eye is less efficient at performing letter identification in bandpass noise. We did not find an overall significant difference with either scrambling type when comparing efficiency between the amblyopic and fellow eye, but we did find such a difference when partitioning our participants by their stereopsis ability. In further analyses of the pattern of mistakes, we found the amblyopic eye shows a distinctive behaviour which correlates with the acuity deficit for both types of scrambling. These results demonstrate that our scrambled stimuli interrogate a component of amblyopic vision that is functionally distinct from that addressed by contrast noise masking.

## 1 Introduction

Amblyopia is a disorder of visual development. It occurs when the brain receives an impoverished input from one eye during childhood. Common causes include visual axis misalignment (strabismus) or unilateral blur (anisometropia). Clinically, it is diagnosed by a reduced optotype letter acuity in the amblyopic eye while the fellow eye exhibits normal acuity even after the amblyogenic factor is corrected. Amblyopia is the most common visual disorder in children, and has a substantial negative impact on quality of life (Birch & Kelly, 2023). The neural basis of the acuity loss in amblyopia remains elusive despite decades of research into the disorder (Kiorpes & McKee, 1999; Levi, 2013). Understanding the sites and mechanisms of this acuity loss will be crucial to developing new treatment options that can recover both monocular acuity and binocular functions (Hess et al., 2014; Hess & Thompson, 2015; Levi, 2020).

Although optotype acuity on the letter chart is the standard clinical measure of visual function, greater insight can be found by measuring the Contrast Sensitivity Function. Measurement of the just-detectable threshold contrast for sinusoidal gratings of different spatial frequencies allows one to define the “window of visibility”: the set of stimuli which the participant is able to see (Campbell, 1983). This includes an estimate of their visual acuity from the highest spatial frequency that is visible at 100% contrast. Indeed, measurements of the contrast sensitivity function from amblyopic participants using their impaired eye typically show a reduction in sensitivity and grating acuity (Bradley & Freeman, 1981; Hess & Howell, 1977; Levi & Harwerth, 1977).

Considering that optotype acuity and the contrast sensitivity function both allow measurement of the monocular deficit in amblyopia, we would reasonably expect a strong relationship between the two. However, an early study showed that some patients have spatial distortions (which would affect letter acuity) in their vision, despite having normal contrast sensitivity (Hess et al., 1978). Using spatial alignment and vernier acuity tasks, participants with strabismic amblyopia show spatial localization deficits that are much worse than what would be predicted based on the reduction in contrast sensitivity alone (Hess & Holliday, 1992; Levi & Klein, 1985). Using a factor analysis on a large sample of amblyopia patients, McKee et al. (2003) identified acuity and contrast sensitivity as separate dimensions on which different clinical subtypes fall.

One explanation for the acuity loss relates to the scrambling found by Hess et al. (1978). Under the spatial scrambling hypothesis the deficit can be attributed, at least in part, to a neural disarray of uncalibrated connections (Hess, 1982; Hess & Field, 1994). This idea has received support from converging lines of evidence, including drawings made by participants (Barrett et al., 2003; Hess et al., 1978; Pugh, 1958), behaviour in psychophysical tasks (Bedell & Flom, 1981; Bedell & Flom, 1983; Fronius & Sireteanu, 1989; Hess & Field, 1994; Lagrèze & Sireteanu, 1991; Piano et al., 2015, 2016; Sireteanu et al., 1993; Wilson, 1991), functional brain imaging (Clavagnier et al., 2015; Li et al., 2007; Szinte et al., 2024), and physiological studies in amblyopic model animals (Bi et al., 2011; Swindale & Mitchell, 1994; Tao et al., 2014). A competing hypothesis attributes the acuity loss to under-sampling. Under that theory, fewer neurons respond to the amblyopic eye, leading to a reduced representation which particularly affects higher spatial frequencies (Levi et al., 1987, 1999; Levi & Klein, 1986, 1996; Sharma et al., 1999).

Psychophysically, one common approach used to tease apart the mechanism underlying a visual function is the noise masking paradigm (Pelli, 1981; Pelli & Farell, 1999). Under the Linear Amplifier Model, the general principle is that the visual system’s performance measured on tasks where stimuli are affected by a known “external” noise can be used to infer the magnitude of that external noise that is equivalent in its effects to the “internal” noise in the system. Also, performance in high levels of external noise can be used to quantify the efficiency with which the system processes that noisy input. In contrast, Perceptual Template Model explicitly assumes both additive and multiplicative noise sources (Lu & Dosher, 1998). Several studies have used these methods to reveal insights into the neural basis of amblyopia. The results and interpretation of these studies depend both on the choice of the external noise that is used (Allard & Faubert, 2014; Baker & Meese, 2012), and the model that is assumed when analysing the data (Baldwin et al., 2016; Lu & Dosher, 1999). Previous studies using white noise and similar broadband noise have argued that the amblyopic eye suffers from a reduced efficiency (Pelli et al. (2004)), an increase in additive internal noise (Levi et al., 2007, 2008; Levi & Klein, 2003; Xu et al., 2006), a combination of the two (Kersten et al. (1988)), or increased multiplicative internal random noise (Levi et al., 2007, 2008).

However, we would not expect the addition of white noise to directly interrogate the spatial disruption that underlies the scrambling hypothesis. This instead calls for a positional noise and a task that relies on positional information (i.e. not the detection of a simple grating). Past studies using this approach have found amblyopic eye performance is limited by its equivalent input noise (not efficiency), for positional hyperacuity (Watt & Hess, 1987) and contour integration (Hess et al., 1997). Since then, similar forms of positional noise have also been applied to investigate the encoding of space in normal subjects (Baldwin et al., 2017; Christensen et al., 2015, 2019) but have not yet been applied in amblyopia.

In this study, we investigate letter identification efficiency for amblyopic and control participants for letters perturbed by positional noise. We chose to use letters to bring our approach into close relation with the letter acuity deficit that is a hallmark of the condition. Moreover, this approach allows us to leverage past studies that investigated the optimal spatial frequency band supporting letter recognition (Chung, Legge, et al., 2002; Chung, Levi, et al., 2002; Majaj et al., 2002; Solomon & Pelli, 1994). This suits the bandpass approach we take to our stimulus design.

To simulate spatial scrambling, we apply a method we recently developed (Zhu et al., 2024). It is inspired by a simple feedforward view of early visual cortex (Hubel & Wiesel (1962)), where neurons in the Lateral Geniculate Nucleus (LGN) first send their projections to simple cells in V1, which then send their own projections elsewhere in the cortex (for example, complex cells or V2) (Alonso et al., 2001). Our approach allows us to simulate spatial scrambling occurring at different stages in this hierarchy.

We refer to a hypothetical scrambling of projections that occurs from LGN to simple cells as **subcortical scrambling** (SCS). Similarly, we refer to the scrambling that occurs in the projections of simple cells as **cortical scrambling** (CS). We mimic the effect of these two types of scrambling by jittering either log-Gabor filters (mimicking oriented receptive fields of simple cells) or their subunits in stimulus space.

There are three main objectives of this study: First, we want to compare the letter identification efficiency for letters affected by both types of scrambling under a common paradigm for both low and medium-to-high spatial frequencies. Second, we want to investigate the link between scrambling and acuity, which we separately measure. Finally, looking beyond efficiency, we want to know if scrambling could alter the pattern of letter classification mistakes. Our results show that the amblyopic eye has normal efficiency for both types of scrambling when we scale by stimulus visibility (Experiment 2A) and acuity (Experiment 2B) despite reduced efficiency for bandpass noise. We find the amblyopic eye makes a distinct pattern of mistakes (Experiment 2A) and the divergence of mistakes between amblyopic and fellow eyes in scrambling noises (Experiment 2B) is correlated with severity of amblyopia (Experiment 1C).

## 2 Experiment 1: Measurements of Visual Acuity

### 2.1 Methods for Experiment 1

We first measured the visual acuity of our participants using three different stimulus designs: grating acuity (**Figure 1**A), broadband letter acuity (**Figure 1**B) and spatially bandpass letter acuity (**Figure 1**C).

**Figure 1:**
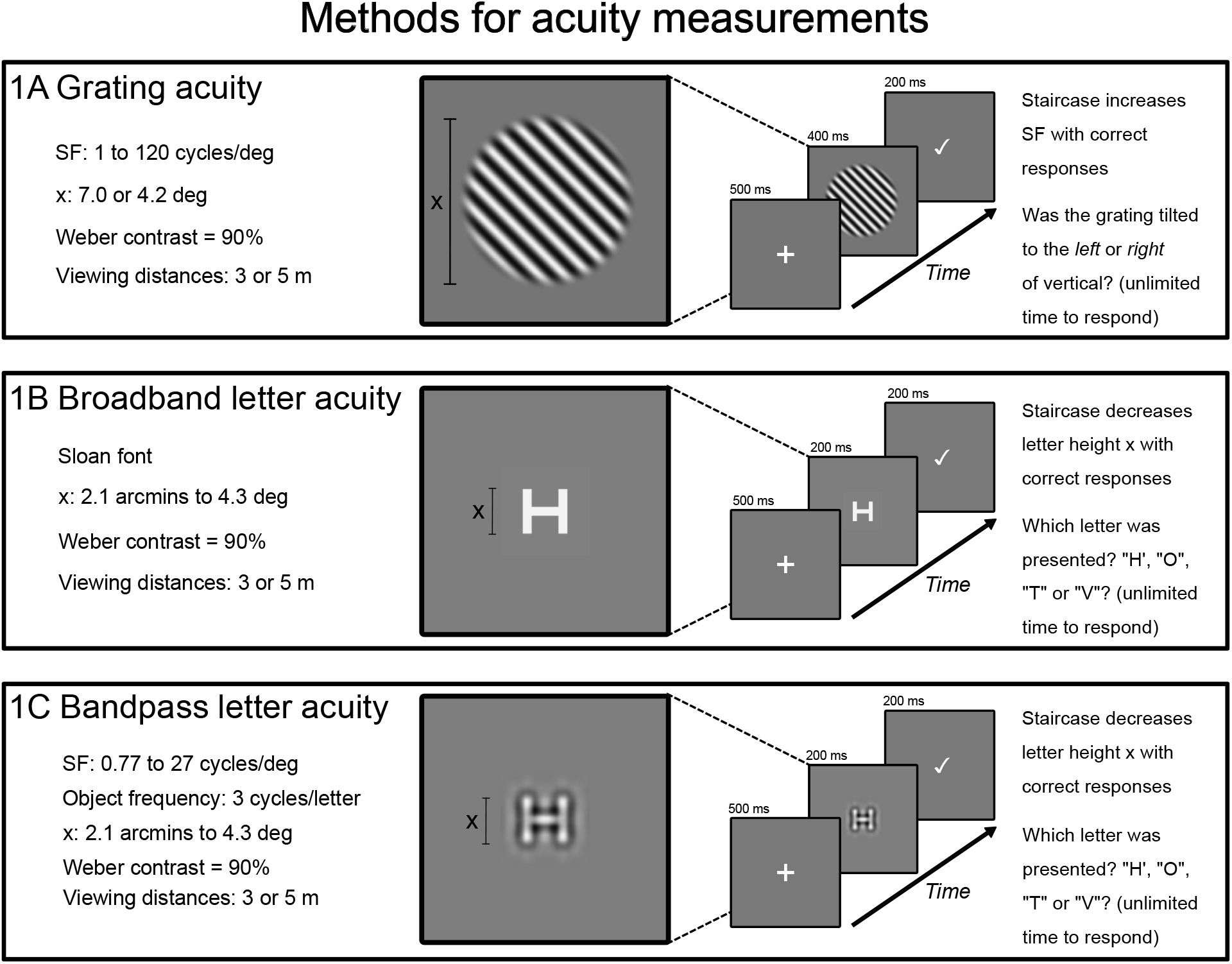
Showing the design of acuity tasks used to quantify the amblyopic deficit.

#### 2.1.1 Participants

We recruited twenty control participants and nineteen amblyopia participants in total. The clinical characteristics of the amblyopic participants are provided in the Appendix (**Table A1**). Twelve control participants (aged 20-30, 7 female, 5 male) and eighteen amblyopic participants (aged 23-73, 10 female, 8 male) participated in Experiments 1A-C. All control participants had normal or corrected-to-normal vision; their eye dominance was determined as their “sighting dominance” using the hole-in-the-card test. Participants gave written informed consent in accordance with the tenets of the Declaration of Helsinki. All study procedures were approved by the Research Ethics Board of the McGill University Health Centre.

#### 2.1.2 Apparatus

Participants viewed a gamma-corrected Display++ LCD screen (Cambridge Research Systems, United Kingdom). The mean luminance of the display was 58 cd/cm^2^. The software ran on a Lenovo P53s laptop (Lenovo Group Limited; Quarry Bay, Hong Kong). The experiment software was written in Python (3.8), making use of the PsychoPy (2021.2.3) library (Peirce, 2007). All experiments were performed indoors under dimmed ambient lighting. For all tasks, monocular viewing was achieved by having the participant wear a black fabric eye patch. Each experiment testing block took between 5 and 10 minutes, with the eye covered by the patch alternating between each testing block. This counter-balanced and helped reduce the potential build-up of adaptation or other effects of monocular deprivation that the patching may have induced (Lunghi et al., 2011; Min et al., 2018; Zhou et al., 2013).

#### 2.1.3 Grating acuity experiment (Experiment 1A)

We generated sinusoidal gratings at two oblique orientations: 45° clockwise or counter-clockwise of vertical. To measure acuity, the spatial frequency of the grating was varied. The absolute size of the stimulus (in degrees of visual angle or “deg”) was constant, meaning that the number of grating cycles in the stimulus scaled linearly with the spatial frequency. Stimuli were presented at 90% Weber contrast.

The threshold spatial frequency at which the grating orientation could be identified was measured using a single-interval two-alternative forced-choice design. Each trial began with a fixation cross that appeared for half a second followed by a 450 ms grating stimulus (FWHM; full-width at half-magnitude of a raised-cosine temporal envelope with a 50 ms ramp and 400 ms plateau). Participants were given unlimited time to respond. Feedback was given at the end of each trial.

Measurements could be made at two viewing distances: 5 m and 3 m. This allowed us to test more severe amblyopic participants who could not identify the orientation of the gratings at 5m through their amblyopic eye. The raised-cosine mask used as a spatial envelope was the same on-screen size regardless of viewing distance. At 5 m, the gratings had a nominal size (full-width at half magnitude) of 3.35 deg. At 3 m this increased to 5.58 deg. Due to the steep fall-off in sensitivity for higher spatial frequencies close to the fovea (Baldwin et al., 2012; Watson, 2018), we expect participants made use of only the very centre of the grating when close to threshold.

Therefore, these differences in stimulus size are immaterial.

The spatial frequency of the grating was controlled by a two-up one-down staircase (terminating after reaching both thirty trials and eight staircase reversals). The starting spatial frequency of the grating was 2.4 cycles/deg at 5 m and 1.4 cycles/deg at 3m. The spatial frequency increased or decreased by a multiplicative factor after each trial, based on whether the response was correct or incorrect. This factor was 1.58x for the first four reversals, and then reduced to 1.26x for the remainder of the block. Data were combined over three repetitions of the task for analysis.

#### 2.1.4 Letter acuity experiments (Experiments 1B and 1C)

Letter acuity was measured using Sloan majuscules (**Figure 2**A-D). We presented single letters in isolation to avoid potential crowding effects (Levi, 2008). As our main results concern the identification of bandpass letters, we used our noiseless letter synthesis algorithm to generate bandpass versions of the Sloan letters (**Figure 2**E-H). This allowed us to measure acuity for both broadband and bandpass letters.

**Figure 2:**
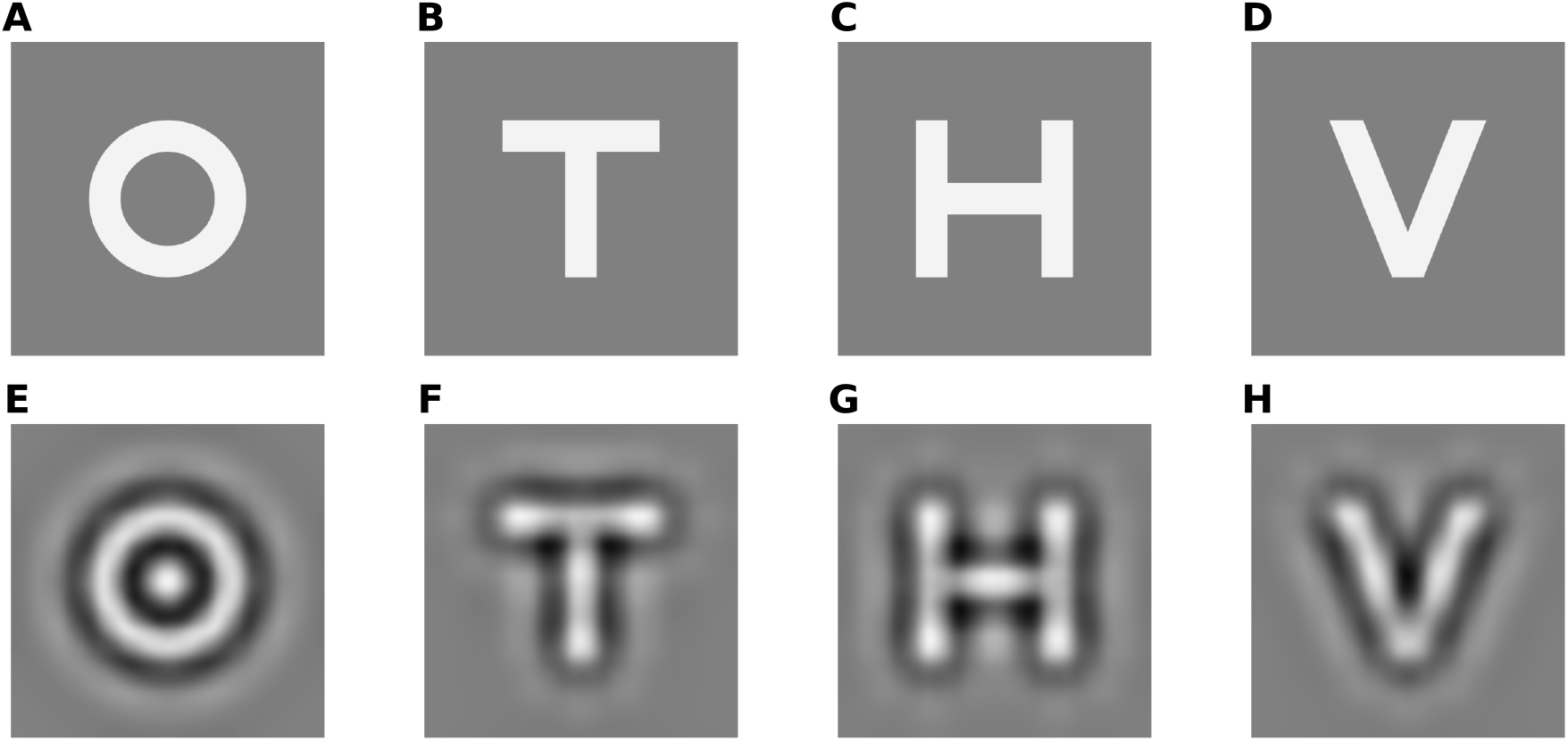
Stimuli used in letter acuity experiments. The top row shows broadband letters. The bottom row shows bandpass letters with their peak spatial frequency determined by Equation 1.

Bandpass letters were filtered using the algorithm described in **Section 3.1.3**, with the optimal filter frequency (f_opt_) in c/deg calculated as

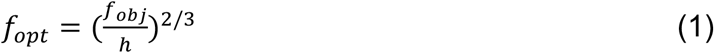

where f_obj_ is the object frequency (for which we chose 3 cycles per object), and h is the letter height in deg. Note that without the exponent of 2/3 Equation 1 would indicate scale invariance. The inclusion of this nonlinear scaling is informed by previous work on letter identification (Chung, Legge, et al., 2002; Majaj et al., 2002).

Both letter acuity tasks used the same basic design as the grating acuity task (1A). The size of the letters was controlled by the staircase, with testing again performed at the two viewing distances as needed. The starting letter height was 2.6 deg at 5 m and 4.4 deg at 3 m. Data were combined over three repetitions of the task as before. To find the threshold spatial frequency (acuity), the letter sizes were converted to their equivalent spatial frequencies using Equation (1) and before psychometric function fitting.

#### 2.1.5 Analysis of Experiment 1A-1C

We fit cumulative normal psychometric functions through Maximum Likelihood Estimation (Kingdom and Prins, 2016), using the scipy.optimize.minimize function from Python’s SciPy package (Virtanen et al., 2020). The slopes and thresholds were fit as free parameters. The lapse rate was fixed at 0.01, and the guess rate was set to either 25% (for 4AFC tasks) or 50% (2AFC). The thresholds were found as the mid-point between guessing and saturated performance (62.5% for 4AFC, 75% for 2AFC).

All statistical analyses were performed in R (R Core Team, 2023). For each experiment, we first removed any extreme outliers defined as being outside the outer fences (3 times the interquartile range) constructed using Tukey’s method (Tukey, 1977). This, in practice, never removed more than two participants in any condition we compared. We performed two-way mixed ANOVA with eye (strong or weak) as a within factor, and group (control or amblyopic) as a between factor, followed by post-hoc paired t-tests or permutation tests. In cases where we compared performance between control and amblyopic groups directly, a regular T-test or Welch T-test was used, depending on the variance assumption. Unless specified, T-tests were always two-tailed. Normality was checked using the Shapiro-Wilk test and a Q-Q plot. Homogeneity of variance was checked using the Levene test. Bonferroni correction for multiple comparisons was used to control false discovery rate. In the results we report the corrected p values. In cases where the correct p value was greater than one we capped the p value at one.

### 2.2 Results of Experiment 1: Measurements with three acuity tasks

The results for acuity experiments (Experiment 1A-C) are shown in **Figure 3**. As expected, comparison of the amblyopic eye against the fellow shows it to have reduced grating acuity (t_18_ = 5.07, p < 0.001), bandpass letter acuity (t_18_ = 5.61, p < 0.001), and broadband letter acuity (t_18_ = 6.15, p < 0.001). Curiously, in control participants we find a slight performance advantage in their non-dominant eye compared to the dominant eye (t_11_ = -3.68, p = 0.008 for bandpass letters; t_11_ = -4.42, p = 0.002 for broadband letters). This was remarkable but not too surprising as sensory dominance does not necessarily align with the sighting dominance we used to classify the control participants’ dominant and non-dominant eyes (Hunter, 2022).

**Figure 3:**
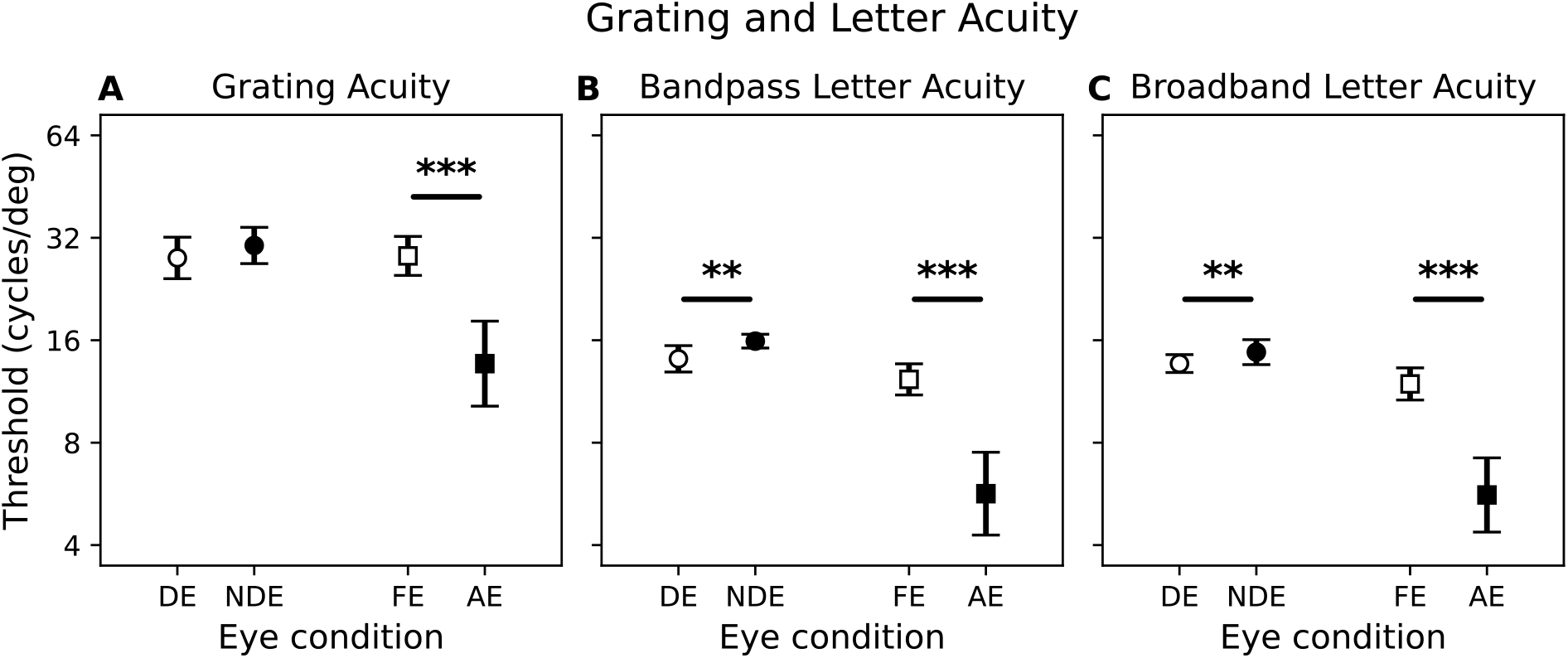
Grating acuity, bandpass letter acuity, and broadband letter acuity. DE: dominant eye, NDE: non-dominant eye, FE: fellow eye, AE: amblyopic eye. Markers represent the mean of the data with error bars indicating the 95% confidence intervals. N=12 for control group and N=18 for the amblyopia group. Significance labels: p<0.05*, p<0.01**, p<0.001***.

In **Figure 4**, we plot the different acuity measures against each other. On double-log axes, the relationships are well-fit by simple linear models (R = 0.88, p < 0.001 for grating against bandpass letter acuity; R = 0.87, p < 0.001 against broadband letter acuity). We find very high agreement between bandpass and broadband letter acuity (R = 0.97, p < 0.001), as predicted by our use of Eq. 1 to select the optimal frequency.

**Figure 4:**
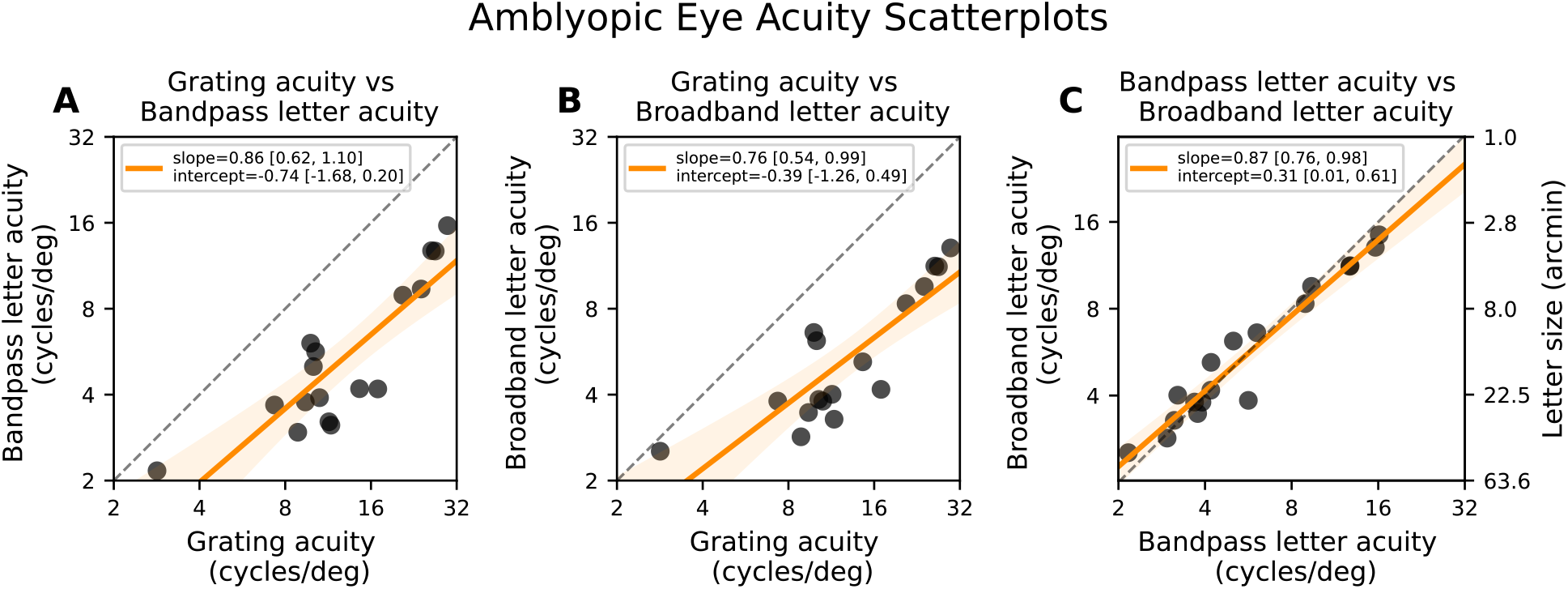
Scatterplots of amblyopic eye acuities against each other. The axes are log-log scaled. The orange line represents the regression line of best fit through the data. The equivalent letter height for the bandpass letter spatial frequency and broadband letter targeted spatial frequency (Equation 1) is shown on a secondary axis on the right. The shaded region represents the 95% confidence interval of the fit. Slope and intercept parameter values of the best fit are shown in the legend along with their 95% confidence intervals.

## 3 Experiment 2: Identification of noisy or scrambled letters

### 3.1 Methods for Experiment 2

We then measured the participants’ ability to identify letters in limiting external noise or scrambling. An overview of the experiments is shown in **Figure 5**. The first variation of this task (Experiment 2A) measured performance for relatively “large” letters at a “low” spatial frequency of 1.5 cycles/deg. We expected performance on this task would be measurable with either eye, even in more severe amblyopic participants. To equate the visibility of the letters in 2A, they were presented at four times their contrast threshold (Hess & Bradley, 1980). Experiment 2B was a second variation of the task, in which we equated visibility of the letters by scaling their size. Using the visual acuity from each eye, the letters were presented at twice the threshold size.

**Figure 5:**
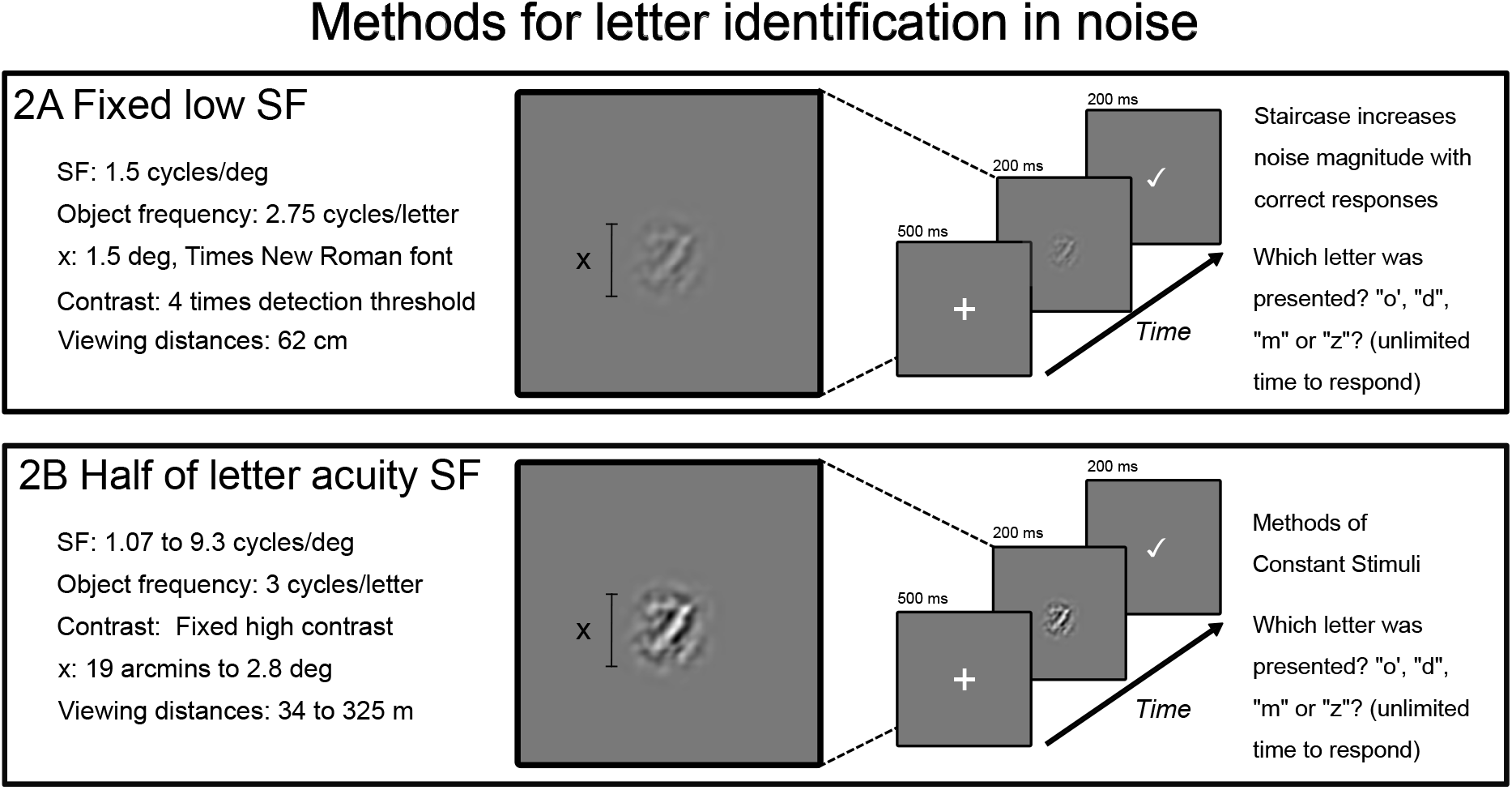
Showing the design of the letter identification in noise tasks.

#### 3.1.1 Participants

Eighteen control participants (aged 19-30, eleven female, seven male; ten of whom also participated in Experiment 1A-C) and eighteen amblyopic participants (aged 23-73, 11 female, 7 male, seventeen of whom also participated in Experiment 1A-C) participated in Experiment 2A. The same twelve control participants and eighteen amblyopic participants that participated in Experiments 1A-C participated in Experiment 2B.

#### 3.1.2 Apparatus

Participants viewed the same gamma-corrected Display++ LCD screen (Cambridge Research Systems, United Kingdom) as in Experiments 1A-C. The experiment software ran on a Dell XPS laptop (Dell Technologies, Inc.; Round Rock, Texas) for experiment 2A. For experiments 2B, the software ran on the same Lenovo P53s laptop that powered the acuity experiments (Lenovo Group Limited; Quarry Bay, Hong Kong).

#### 3.1.3 Stimulus synthesis algorithm

For our letter identification experiments, we used four lowercase letters from the Times New Roman font: o, d, m, z. These were chosen as they are perceptually distinct from each other (Courrieu et al., 2004; Janini et al., 2022). Our stimulus generation algorithm (described below) effectively bandpass-filtered the letters to target a spatial frequency channel that should be optimal for letter identification. Based on previous findings with similar fonts, we selected an object frequency of 2.75 cycles per letter for Experiment 2A and 3 cycles per letter for Experiment 2B (Chung, Legge, et al., 2002; Chung, Levi, et al., 2002; Majaj et al., 2002; Pelli et al., 2004; Solomon & Pelli, 1994).

The method used to generate stimuli was based on a wavelet transform using log Gabor filters (Fischer et al., 2007; Würtz, 1995). We chose log-Gabor filters because they approximate simple cell receptive field properties, having symmetrical tuning on a log-frequency axis with the additional benefit of being DC-balanced in any phase (Daugman, 1985; Field, 1987). The log-Gabors we used are Cartesian-separable, generated (for a full description see Baker et al., 2022; Meese, 2010).

To synthesise our letter stimuli, we began with a bitmap image with the letter rendered in the centre. This image was convolved with a bank of 2D log-Gabor filters to perform a decomposition into frequency and orientation sub-bands. All filters had a single peak spatial frequency, with a bandwidth of 1.5 octaves. In orientation, the filters had a bandwidth of 30°, with eight peak orientations evenly spaced between 0 and 360° (0, 45, 90, 135, 180, 225, 270, 315). For each orientation we used a quadrature pair of filters, with phases at 0° and 90°. This results in a total of 16 log-Gabor filters (in each experiment). Decomposition of a letter image therefore gave us 16 spatial maps of filter weights (one map per filter).

The maps of filter weights were then downsampled at a sampling grain of every half-wavelength to avoid redundancy of coding for nearby positions (Field, 1987). We applied a threshold to keep the top 1% of most active weights across the 16 maps. In the resynthesis, we convolved each one of these maps with the corresponding log-Gabor wavelet to render 16 images, each one contributing wavelet components of a specific spatial frequency, orientation, and phase. These were then linearly summed into a reconstructed image which is normalised to a specified RMS contrast. For our letters without scrambling, this completes the process (**Figure 6**, first column).

**Figure 6:**
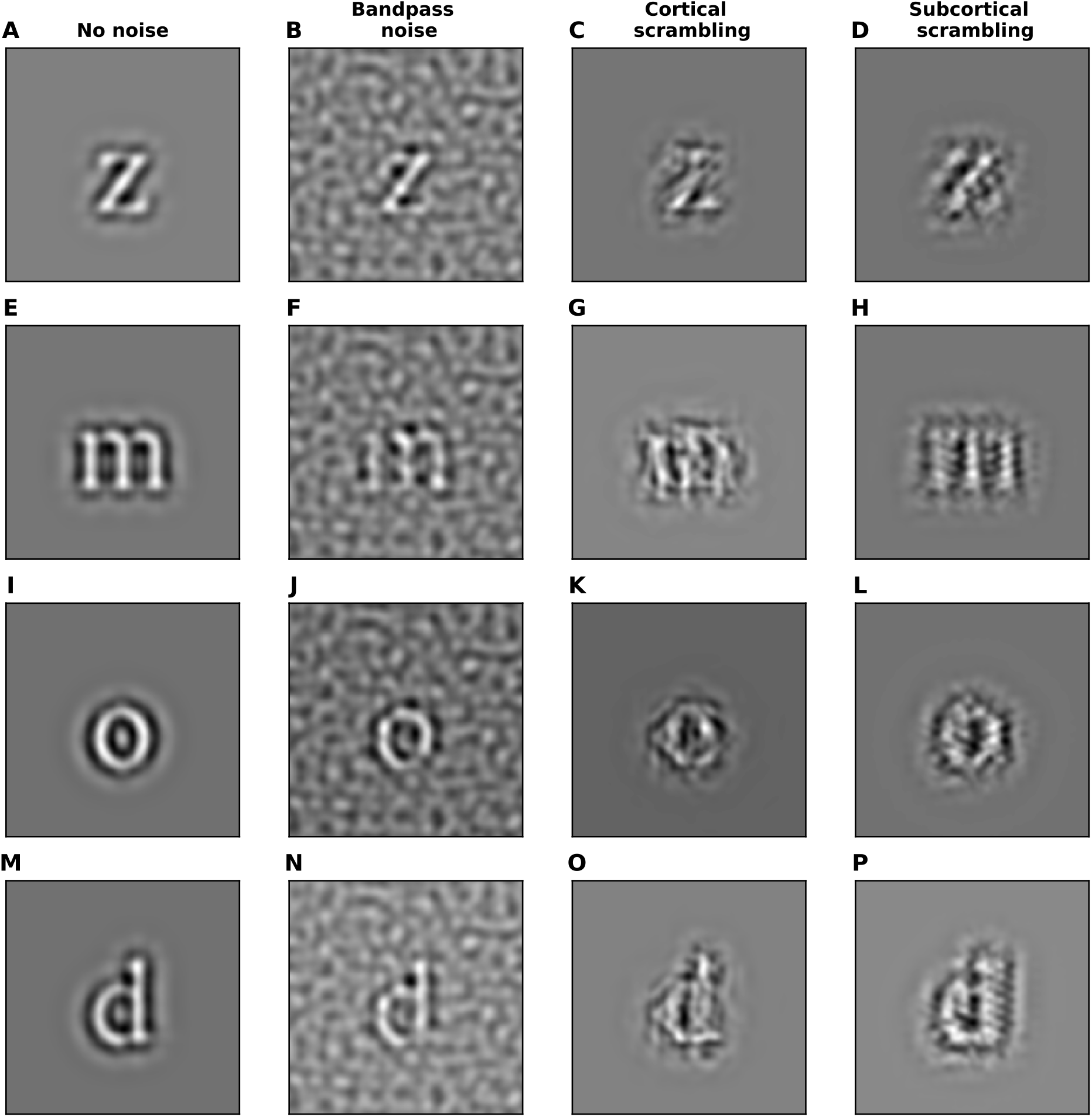
The four columns (in order) show letters: i) without noise or scrambling, ii) embedded in bandpass noise, iii) affected by our “cortical” scrambling algorithm, and iv) affected by our “subcortical” scrambling algorithm. The magnitude of noise and scrambling levels are set to half the average threshold measured in Experiment 2A.

For the bandpass noise (BN) condition, we fed white noise through the same process used to synthesise our letter stimuli. The output was added to the bandpass letter before then normalising the RMS contrast (**Figure 6**, second column). Adding different magnitudes of noise therefore results in a stimulus of the same overall contrast, but with that contrast being contributed by different proportions of signal (letter) and noise.

For the cortical scrambling (CS) condition, we jittered the positions of the weights (representing local oriented features of a particular phase and spatial frequency) in the 16 spatial maps before resynthesis. The translation applied to each weight was independently drawn from a 2D Gaussian probability distribution. The standard deviation (with the same value used for the vertical and horizontal spread) determined the magnitude of the cortical scrambling noise (**Figure 6**, third column).

For subcortical scrambling (SCS), we simulate scrambling within the local feature detectors that are modelled by our log-Gabor wavelets. This is accomplished by deconvolving the log-Gabor with an isotropic log-Gabor kernel. This created a “wiring diagram” that describes the spatial arrangement of non-oriented subunits that would form the oriented log-Gabor, analogous to models of simple cell receptive field formation from LGN afferents (Alonso et al., 2001; Hubel & Wiesel, 1962). The subcortical scrambling is modelled as jitter applied to the locations of the inputs in that wiring diagram. Convolving the scrambled wiring diagram with the isotropic log-Gabor kernel creates a scrambled wavelet. To create our SCS letter stimuli, we generate these scrambled wavelets (one for each of the 16 combinations of orientation and phase) and use those in place of the original wavelet when synthesising the stimulus (**Figure 6**, rightmost column). The effects of SCS on the spatial and Fourier profiles of an individual log-Gabor wavelet are visualized in **Figure 7**. We see that SCS spreads energy into neighbouring orientations while keeping within the same spatial frequency band.

**Figure 7:**
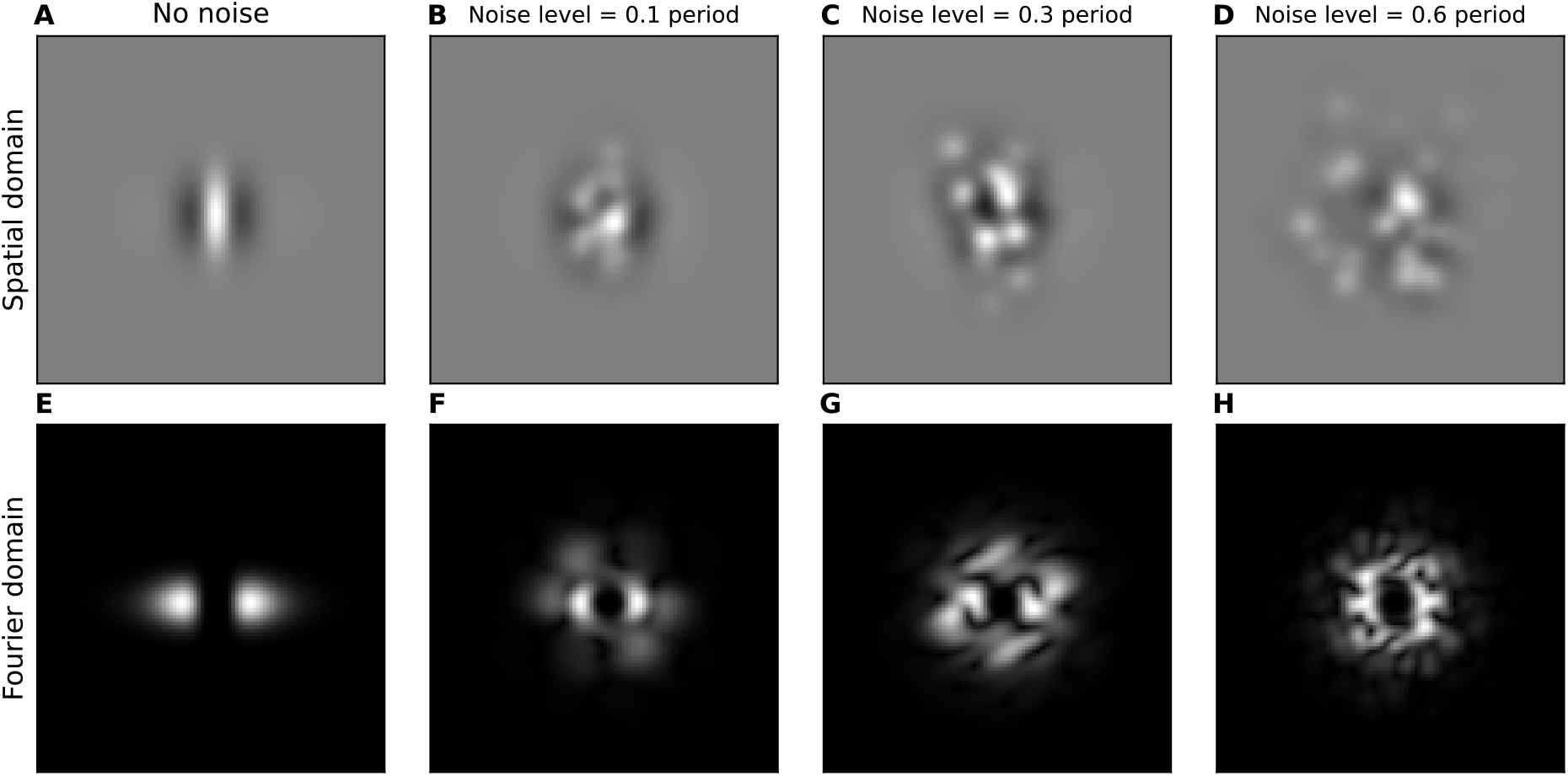
The effect of subcortical scrambling in spatial (top) and Fourier (bottom) domains.

#### 3.1.4 Letter identification in noise at a low spatial frequency (Experiment 2A)

A reduction in contrast sensitivity in the amblyopic eye is a hallmark feature of the amblyopic phenotype (Bradley & Freeman, 1981; Hess & Howell, 1977; Levi & Harwerth, 1982). The scrambling we wish to investigate is an additional separable amblyopic deficit which may be profound even when contrast sensitivity for detection is relatively unaffected (Hess, 1982). We set out to measure scrambling independent of contrast sensitivity by normalising the stimuli shown to each eye to be presented at contrasts which were above that eye’s detection threshold by the same factor. For this reason, we began our testing by measuring the contrast threshold for identifying (noiseless) bandpass letters.

We used the same basic task design as in Experiment 1. The viewing distance for both tasks was 62 cm. At this distance, there are 31 pixels/deg. The staircase controlled the contrast of the letter stimulus, with an initial Weber contrast of 7%. This was equivalent to 0.01 RMS contrast, defined as the standard deviation of the luminance across the stimulus region. Contrast thresholds (**Table 1**) were obtained by fitting a psychometric function to the data from three repetitions of the task.

**Table 1:**
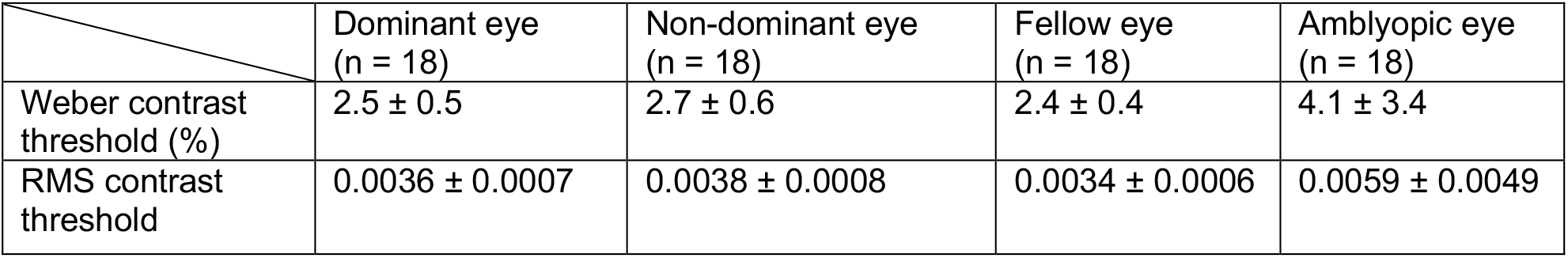
Contrast thresholds (given as both Weber and equivalent RMS contrast) for noiseless letter identification. Reported values are the mean and standard deviation of contrast thresholds across all participants.

|  | Dominant eye<br>(n = 18) | Non-dominant eye<br>(n = 18) | Fellow eye<br>(n = 18) | Amblyopic eye<br>(n = 18) |
| --- | --- | --- | --- | --- |
| Weber contrast threshold (%) | 2.5 ± 0.5 | 2.7 ± 0.6 | 2.4 ± 0.4 | 4.1 ± 3.4 |
| RMS contrast threshold | 0.0036 ± 0.0007 | 0.0038 ± 0.0008 | 0.0034 ± 0.0006 | 0.0059 ± 0.0049 |

In our main task, the stimulus contrast was fixed at four times the threshold RMS contrast. We then measured the threshold noise or scrambling magnitude for letter identification. Because the RMS contrast reflects the square root of the contrast energy of the stimulus, whereas the Weber contrast measures the peak deviation from the mean luminance, the two values had only an *expected* scaling factor that could be used to translate one to the other for our stochastically generated stimuli. The expected RMS and Weber contrast for the different noise and viewing conditions is shown in **Table 2**. As a greater noise or scrambling magnitude made the task more difficult, we flipped the staircase such that correct responses increased the noise magnitude and incorrect responses decreased it. The starting value was 0.1 (the reciprocal of signal-to-noise ratio calculated in RMS contrast unit) for the BN, 4 arcminutes for SCS and CS. Data for three repetitions of the task were collected, with thresholds obtained by fitting a psychometric function to the combined data.

**Table 2:** RMS contrast and expected Weber contrast of stimuli (%) in the scrambled letter identification for low spatial frequency experiment. Reported values are the mean and standard deviation of Weber contrasts across all noise levels used for a given noise condition

|  | Dominant eye | Non-dominant eye | Fellow eye | Amblyopic eye |
| --- | --- | --- | --- | --- |
| RMS Contrast (n = 18) | 0.014 ± 0.003 | 0.015 ± 0.003 | 0.014 ± 0.003 | 0.024 ± 0.020 |
| Bandpass noise | 7 ± 2 | 8 ± 2 | 7 ± 2 | 12 ± 3 |
| Cortical scrambling | 11 ± 2 | 12 ± 2 | 10 ± 2 | 18 ± 3 |
| Subcortical scrambling | 10 ± 2 | 11 ± 2 | 10 ± 2 | 16 ± 3 |

#### 3.1.5 Letter identification in noise for mid-high spatial frequencies (Experiment 2B)

Where Experiment 2A tested participants’ ability to identify letters at a single, lower spatial frequency (1.5 cycles/deg) at which the task was possible for our control and amblyopic participants using either eye, Experiment 2B sought to measure performance closer to the acuity limit for each eye tested. We used the same 4AFC design and temporal profile as in Experiment 1. However, to allow for a double-pass analysis we switched from a staircase design to a method of constant stimuli. We sampled 20 trials (5 of each of our 4 target letters) at five noise magnitudes for a total of 100 trials per fitted psychometric function. The same set of stimuli were shown to each participant in two repetitions, with the order randomised. Thresholds were derived from fitting the combined data across the two repeats of the experiment.

In Experiment 2B, we normalised letters to be equally discriminable by their size, rather than the contrast normalisation used in Experiment 2A. We filtered the image with an object frequency of 3 cycles/letter. The viewing distance for each eye was varied so that the spatial frequency channel targeted was half the bandpass letter acuity (Experiment 1C). Viewing distance ranged over a factor of ten (from 0.3 m for the amblyopic participant with the worst acuity, to 3.3 m for the most keen-eyed control participant).

Stimuli were shown at a common RMS contrast, calculated to be the maximum that could be rendered while avoiding “clipping” from attempting to show contrasts beyond the range of the display. The expected Weber contrast of our noisy stimuli was 82% for the BN condition, 90% for CS, and 85% for SCS.

#### 3.1.6 Analysis of Experiment 2A-2B

To compare performance, we extracted thresholds using same psychometric function fitting methods as the acuity experiments, fitting a cumulative normal function to participant data. All statistical testing was carried out as described in Section 2.1.5.

To compare mistake patterns, we constructed confusion matrices that capture the pattern of classification responses across all noise levels tested (see **Appendix A2** for an example). The four diagonal entries of the matrix are counts of correct letter classification (i.e. “shown m, responded m”). The off-diagonal elements are counts of mistakes. By removing the correct responses, an analysis of *just* the mistakes provides independent information about the classification behaviour.

To compare the mistake similarity between eyes within each participant, we make use of the Kullback–Leibler (KL) divergence (Kullback & Leibler, 1951). For each target letter, we are interested in how much each eye’s distribution of mistakes differs from the pooled distribution of the mistakes from both eyes. We quantify this as the KL divergence between the probability distributions. Then sum the KL divergences, with each weighted by the number of trials from which it is calculated

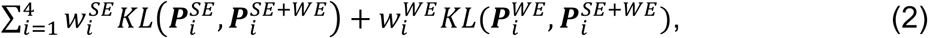

where *i* indexes the target letter, 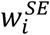 and 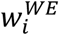 are the weight for KL divergence for the strong and weak eyes respectively, 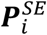 and 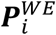 are normalized probability vectors (of length 3) for mistake frequencies of strong eye and weak eye, and 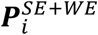 is the normalized probability vector (of length 3) for mistake frequencies of the combined data of strong and weak eye.

KL divergence (Kullback & Leibler, 1951) for a discrete probability distribution is defined as:

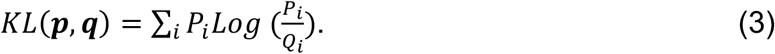

Our metric indicates the dissimilarity of either eyes’ mistake patterns from the average mistake pattern of both eyes. This approach provides two advantages compared with calculating KL divergence between strong and weak eyes directly: i) our metric is symmetric, and ii) our metric gracefully handles edge cases in our small sample data where a mistake occurs in one eye (a non-zero value in *p*_*i*_) that never occurs in the other (a probability of zero in *Q*_*i*_) The KL divergence in that case would be undefined if we were simply comparing one eye to the other. Because we pool the data from the two eyes in *Q*_*i*_, there can never be an event in *p*_*i*_ that has zero probability in *Q*_*i*_.

Beyond comparing mistakes between eyes within a single individual, one can also make group-level comparisons. We compared the mistakes of each of four groups of eyes, derived from the two eyes of our two participant groups. These were the amblyopic and fellow eyes of our amblyopic participants, and the non-dominant and dominant eyes of our control participants. For this analysis we chose to use the Deviance rather than the KL divergence to compare the mistake distributions. The deviance is defined as:

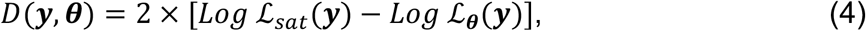

where *y* are the data, *θ* represent the model parameters, *Log* ℒ_sat_(*y*)is the Log likelihood of the data under the saturated model that generates predictions with the same probability distribution as *y, Log* ℒ_*θ*_(*y*) is the Log likelihood of the data under the model which is being compared. From inspection, one can see that the KL divergence (Eq. 3) and Deviance (Eq. 4) are related. For our use, the difference between our analyses with the KL divergence and with the Deviance is that the latter implicitly weights the values from each participant by the number of trials on which they are based (see **Appendix A3**).

For each eye from each participant, we were interested in how similar its mistake pattern was to that from those four eye groups (amblyopic, fellow, non-dominant, or dominant) from *all other* participants. To normalise this measure, we looked for a similarity to a specific group that was greater than that to the distribution of mistakes from *all* eyes. The deviance is a measure of *dissimilarity*, so we calculate

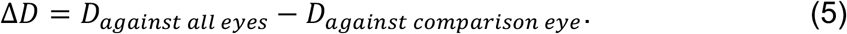

*D*_*against all eyes*_ is defined as:

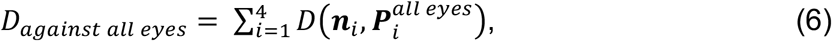

where *i* indexes the target letter, *n*_*i*_ is the mistake count vector (of length 3) for the eye, 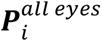 is the normalized probability vector of length 3 for mistakes of combined data across all eyes from all other participants. We define *D*_*against all eyes*_ as

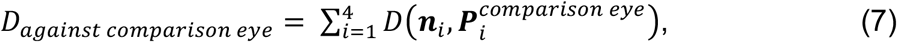

where 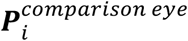 is the normalized probability vector of length 3 for mistakes of the combined data for the comparison eye group from all other participants.

### 3.2 Results of Experiment 2: Identification of noisy or scrambled letters

#### 3.2.1 Identification of 1.5 c/deg bandpass letters in limiting noise or scrambling

The results from Experiment 2A, where the bandpass letters had a spatial frequency of 1.5 c/deg, are shown in **Figure 8**. The contrast of the stimuli was normalised (set to four times the detection threshold) for each eye. Therefore, the comparisons between our amblyopic and control participants, and between the two eyes of each participant group, reflect differences beyond a variation in contrast sensitivity.

**Figure 8:**
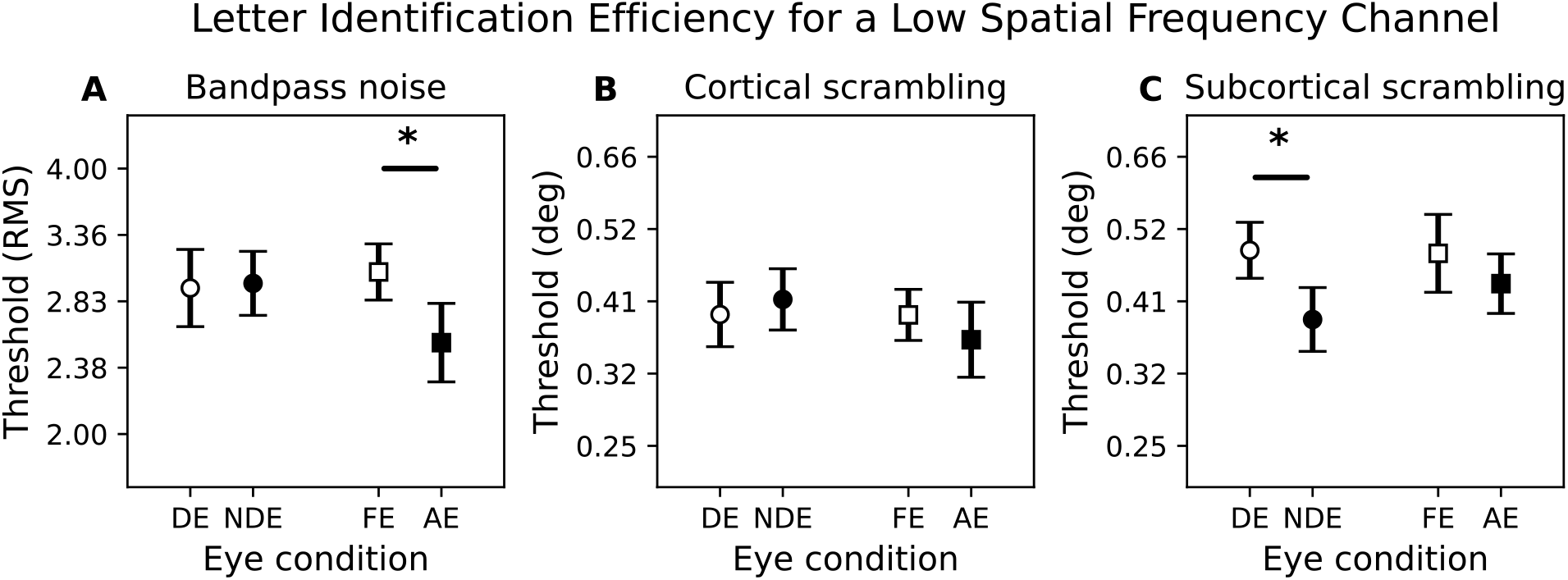
Thresholds for 1.5 cycles/deg channel. DE: dominant eye, NDE: non-dominant eye, FE: fellow eye, AE: amblyopic eye. Markers represent the mean of the data with error bars indicating the 95% confidence intervals. N=18 for both control and amblyopia groups.

For bandpass noise (BN; **Figure 8**A), the relative noise contrast at which the letter could still be identified is lower in the amblyopic eyes of our amblyopic participants. This means that letter identification using those eyes was more susceptible to interference from the masking noise (i.e. the processing is less efficient). This finding is supported by an ANOVA, which reveals a significant main effect of eye (F_1,34_ = 4.38, p = 0.044, generalized eta squared (ges) = 0.046) and a significant interaction between group and eye (F_1,34_ = 5.71, p = 0.023, ges = 0.059). Post-hoc comparisons indicate that within the amblyopic group, threshold in the amblyopic eye (AE) is significantly lower than the fellow eye (FE) (t_17_=2.84, p = 0.022), while no significant difference (t_17_=1.62, p = 1) is observed in the control group between the dominant eye (DE) and non-dominant eye (NDE).

For cortical scrambling (CS; **Figure 8**B), the threshold scrambling magnitude for letter identification appears to be similar across the different groups and tested eye conditions. This is supported by our ANOVA, which reveals no significant main effect (F_1,34_ = 1.12, p = 0.30, ges = 0.021 for group; F_1,34_ = 0.13, p = 0.72, ges = 0.001 for eye) or interaction (F_1,34_ = 2.15, p = 0.15, ges = 0.021).

For subcortical scrambling (SCS; **Figure 8**C), the threshold scrambling magnitude is higher for stronger eye than the weaker eye. This is supported by an ANOVA, which reveals a significant main effect of eye (F_1,34_ = 8.80, p = 0.0050, ges = 0.11). Post-hoc comparisons found this is mainly driven by the control group, in which the threshold for DE is significantly higher than NDE (t_17_=2.66, p = 0.032). No significant difference is observed within the amblyopic group between FE and AE (t_17_=1.43, p = 0.34).

#### 3.2.2 Identification of letters having a spatial frequency that is half the acuity threshold

The second version of our letter identification task was designed to address an amblyopic deficit that is scale-dependent, with the effects restricted to spatial frequencies closer to the acuity limit (Hess et al., 1978). Comparisons at a lower spatial frequency, such as the 1.5 c/deg used in Experiment 2A, may fail to show in amblyopic participants whose vision is unaffected at lower spatial frequencies.

If the difference between the amblyopic eye and fellow eye were simply due to a global loss of sensitivity or lower cut-off spatial frequency then might normalise performance by re-scaling our stimuli to a size that should be equally identifiable (Chung, Levi, et al., 2002; Hess & Holliday, 1992; Levi & Klein, 1982). In this experiment we did so by presenting letter stimuli scaled to be twice the “bandpass letter acuity” size measured in Experiment 1C. The scale was set independently for each eye. These results therefore show effects of amblyopia beyond those attributable to a change in spatial scale.

The noise threshold results are shown in **Figure 9**. For BN (**Figure 9**A), the amblyopic eye threshold is lower than the fellow eye in our amblyopic group (t_17_=3.48, p = 0.0060), while no significant difference is observed within the control group between the non-dominant eye and the dominant eye (t_10_=-0.47, p = 0.65). To compare the scrambling magnitudes (**Figure 9**B-C) between eyes where stimuli were presented at different spatial frequency, we multiply the absolute magnitude by the inverse of the spatial frequency. This gives scrambling in units of “cycles” of a periodic stimulus having the same spatial frequency as the target letter. For CS or SCS, we find no statistically significant difference in scrambling thresholds between eye or group. ANOVA for CS that reveals no significant main effect (F_1,28_ = 1.32, p = 0.26, ges = 0.037 for group; F_1,28_ = 1.10, p = 0.30, ges = 0.007 for eye) or interaction (F_1,28_ = 3.39, p = 0.08, ges = 0.022). ANOVA for SCS also shows no significant main effect (F_1,28_ = 0.44, p = 0.51, ges = 0.012 for group; F_1,28_ = 0.67, p = 0.42, ges = 0.005 for eye) or interaction (F_1,28_ = 1.83, p = 0.19, ges = 0.014).

**Figure 9:**
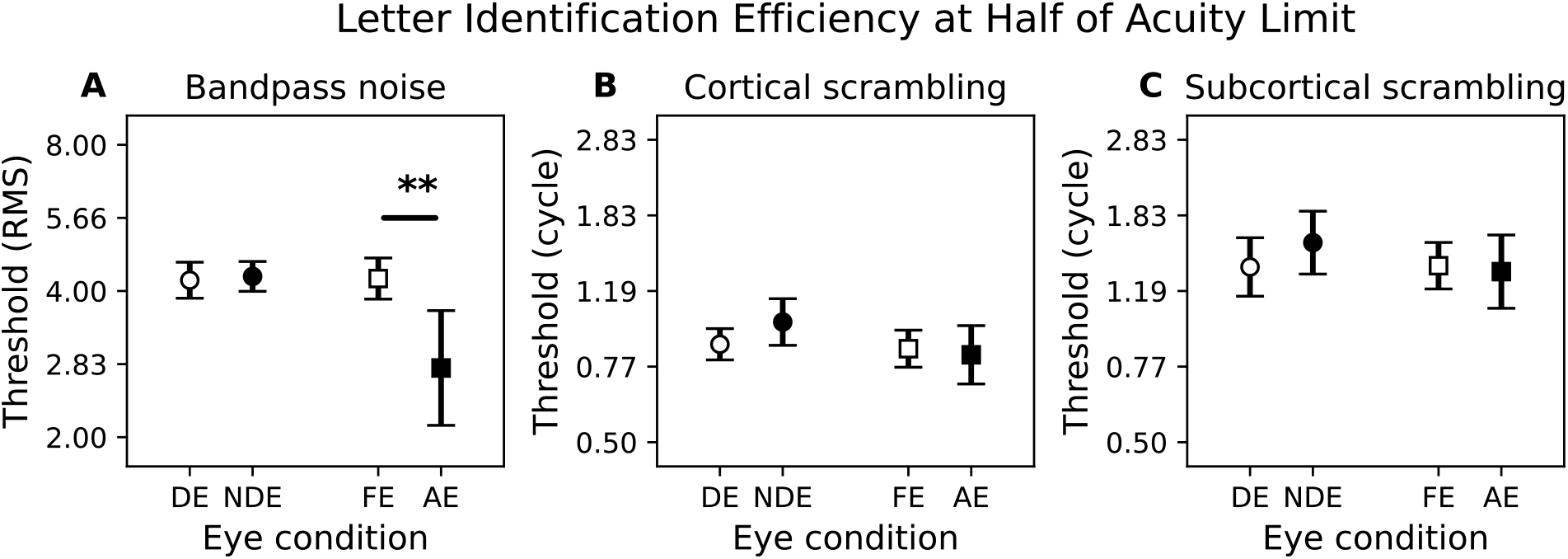
Thresholds for 0.5x letter acuity channel. DE: dominant eye, NDE: non-dominant eye, FE: fellow eye, AE: amblyopic eye. Markers represent the mean of the data with error bars indicating the 95% confidence intervals. N=12 for control group and N=18 for amblyopia group.

#### 3.2.3 The relationship between letter identification thresholds and stereo ability

It has been suggested that the stereopsis deficit found in amblyopia may be due to spatial scrambling (Alarcon Carrillo et al., 2023; Piano et al., 2015). We used the Randot stereo acuity measurements to divide participants into those with “good” stereopsis (< 200 arc sec) and those with poor stereopsis (including no stereopsis).

For both Experiment 1 and 2, we compared the amblyopic eye’s threshold between the two groups (**Figure 10**). In Experiment 1 (**Figure 10** A-C), the threshold is significantly higher (indicating more efficient processing) in the good stereopsis group than the poor/no stereopsis group for both BN (t_16_ = -2.49, p = 0.012) and CS (t_16_ = -2.19,p = 0.022). In Experiment 2 (**Figure 10** D-F), this effect is found for all three noise and scrambling conditions (BN: t_16_ = -2.60,p = 0.0096; CS: t_16_ = -2.51, p = 0.012; SCS: t_16_ = - 2.33, p = 0.018).

**Figure 10:**
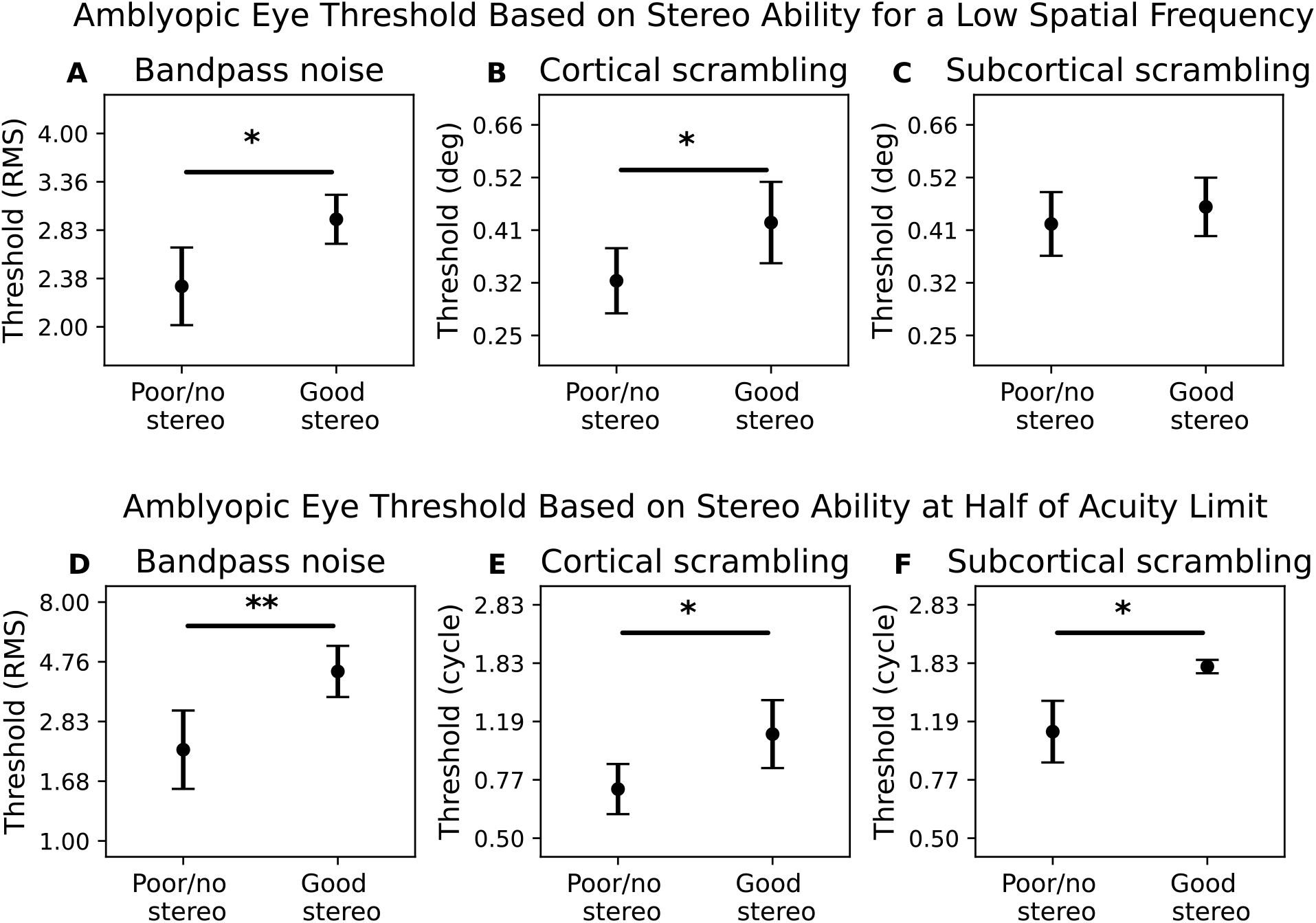
Amblyopic eye noise threshold comparison between participants with poor/no stereopsis and those with “good” stereopsis (Randot stereoacuity £ 200 arc sec). Points represent the mean of the data with error bars indicating the 95% confidence intervals. Top row: N=11 for no stereo group and N=7 for some stereo group. Bottom row: N=12 for no stereo group and N=6 for some stereo group.

#### 3.2.4 Double-pass analysis and comparison of letter mistake patterns

By comparing the “threshold” magnitudes of noise or scrambling at which the task could be performed, the results above indicate the relative efficiency of the letter identification process. The method of constant stimuli design we used in Experiment 2B allowed us to go further, by comparing the responses from two repetitions of the experiment with the same stimuli. We used this to analyse the consistency of the participants’ responses (Burgess & Colborne, 1988; Gold et al., 1999; Green, 1964; Levi et al., 2005, 2007; Levi & Klein, 2003). This quantifies the extent to which the random properties of our noise and scrambling stimuli determine the trial-by-trial responses (including the mistakes). Our analysis (provided in **Appendix A4**) show a significant stimulus-driven factor in letter responses given in trials where participants made a mistake. We also see that our three stimulus conditions lead to different levels of consistency across our participant groups and eye conditions.

Beyond this simple repetition-to-repetition consistency, we hypothesised that distortion due to amblyopic eye scrambling would lead to a distinct *pattern* of letter pair confusions. In an additional analysis, we therefore compared the pattern of mistakes made by our different participant groups using each of their eyes. The metrics we use to compare these mistake patterns are described in the **Methods**. They allow us to analyse the probability distribution of the different mistakes, even in Experiment 2A where the stimuli were not paired between the two repetitions.

We conducted a group-level analysis on the data from Experiments 2A and 2B, in each case we asked if the data from each eye of each participant group was better modelled by the data from *that* eye from all other participants? Or, was its pattern of mistakes similar to the general pattern of mistakes that were found regardless of eye or participant group. For Experiment 2B we found no significant pattern in these results. For Experiment 2A, however, there was a distinctive pattern which we show in **Figure 11**. Each point represents a difference in the Deviance when comparing the data from one eye group (fellow, amblyopic, dominant, or non-dominant) against either the pooled data from all other participants for each eye group (comparisons against each being shown in the north, south, east, and west regions on each plot) or the pooled data from all eye groups from all other participants. A score of zero (the thick line circle) indicates that the deviance is the same for the specific eye group as it is for the data pooled from all eye groups. A positive difference (placing the point outside the circle) means the data more closely resemble that from the comparison eye group. It is important to note that the deviance score is always calculated between independent data, as the pooled data for both all eyes and comparison eyes always exclude the individual eye data.

**Figure 11:**
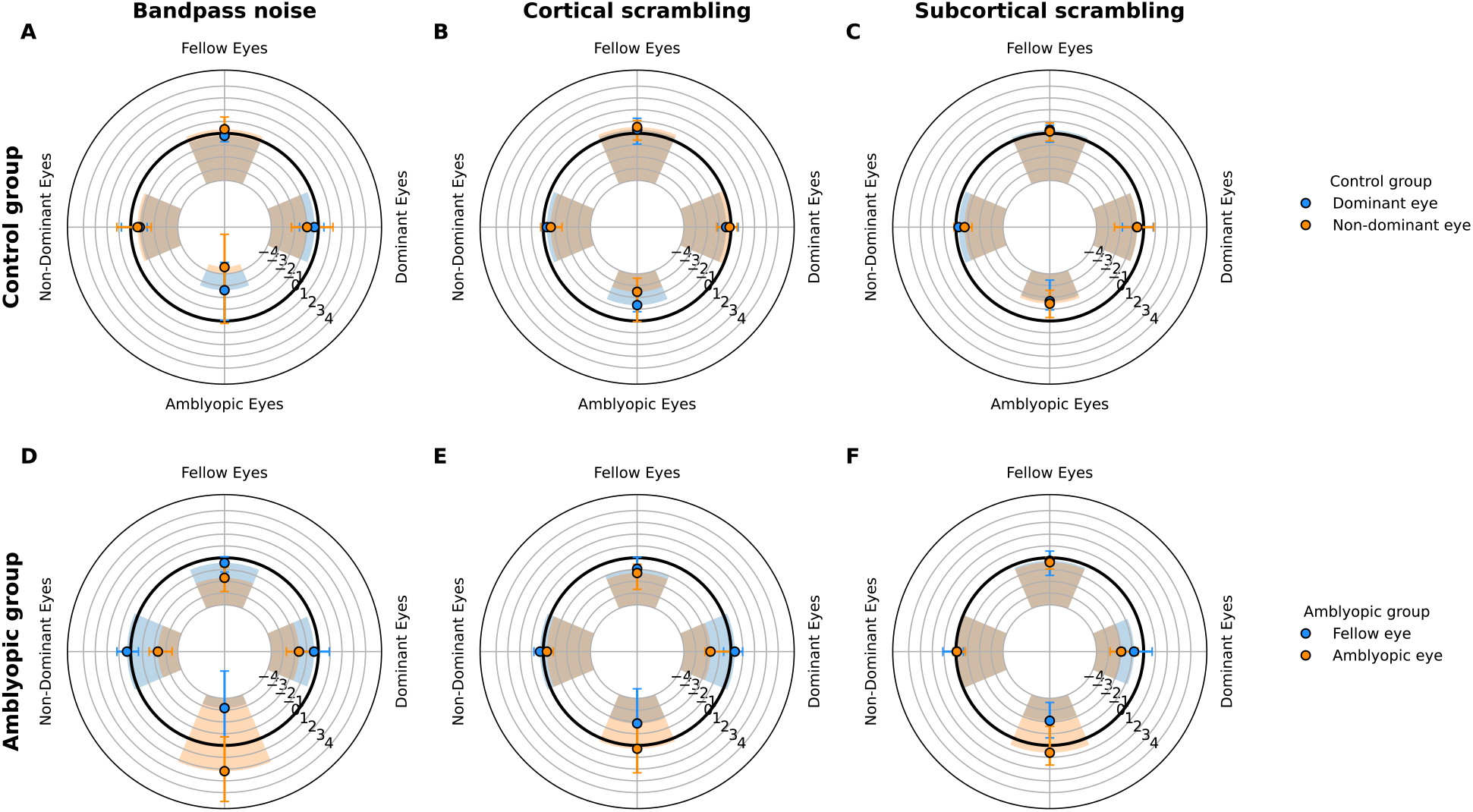
Median similarity of the mistakes made (in Experiment 2A) by each eye compared to the pooled distribution of mistakes from all other participants. Similarity is calculated as the difference in Deviance from comparisons against either i) all eyes, or ii) the comparison group of eyes (labelled at the four cardinal points on each subplot). The two rows show data from 18 control (top) and 18 amblyopic (bottom) participants. The columns show data from the bandpass noise (left), cortical scrambling (middle), and subcortical scrambling (right) conditions. For each group, the blue markers represent the stronger eye (i.e. dominant eye for controls; fellow eye for amblyopic group). The orange markers represent the weaker eye (non-dominant eye for controls; amblyopic eye for amblyopic group). Error bars indicate the 95% confidence interval.

In bandpass noise (**Figure 11**, left column), the amblyopic eye shows a distinct pattern of mistakes. For the comparison against the pooled amblyopic eye data (bottom of each circular plot), we see the amblyopic fellow eye (blue point, bottom plot), and control dominant and non-dominant eyes (blue and orange points, top plot) are all less similar to the pooled amblyopic data than the data pooled over all eyes (FE: t_17_ = -3.43, p = 0.013; DE: t_17_ = -2.95, p = 0.036; NDE: t_17_ = -3.47, p = 0.012). Conversely, when the data from amblyopic eyes (orange points, bottom plot) are compared against the other groups, we find it is less similar to the data from all other pooled eye groups (FEs: t_17_ = - 3.60, p = 0.009; DEs: p = 0.038, permutation test, NDEs: t_17_ = -4.51, p=0.001) and more similar to the pooled amblyopic eye data from other participants. This is supported by an ANOVA, which reveals a significant interaction between group and eye (F_1,34_ = 12.70, p = 0.001, ges = 0.091), with post-hoc comparisons showing a significant difference between fellow and amblyopic (p < 0.001, Wilcoxon signed-rank test).

The two scrambling conditions also show a significant (though less strong) difference in the pattern of mistakes through the amblyopic eye. In cortical scrambling (**Figure 11**, middle column), we find the mistake patterns from FE and DE are less similar to AEs than to the pooled mistakes over all eyes (bottom, FE: t_17_ = -3.52, p = 0.011; DE: t_17_ = - 4.25, p = 0.0022). This is partially mirrored in the comparison against FEs (top) and DEs (right), where we find AE is less similar to FEs (p<0.001, permutation test) but not significantly less similar to DEs (t_17_ = -2.58, p = 0.079). Here we also find that AE is more similar to other AEs than FE. This is reinforced by an ANOVA, with a significant interaction between group and eye (F_1,34_ = 4.96, p = 0.033, ges = 0.049). Post hoc comparisons show a significant difference between FE and AE (t_17_ = -3.62, p = 0.0042).

In subcortical scrambling (**Figure 11** C,F), we find FE and DE are less similar to AEs than all eyes (bottom, FE: t_17_ = -2.93, p = 0.037; DE: t_17_ = -2.94, p = 0.036). However, this is not mirrored when compare AE against FEs and DEs, in which we find AE is not statistically less similar to them (FEs: t_17_ = -1.74, p = 0.40; DEs: t_17_ = -2.36, p = 0.12). We also performed ANOVA for the comparison against AEs and found only a main effect for eye with no interaction (F_1,34_ = 5.54, p = 0.024, ges = 0.061 for eye, F_1,34_ = 1.45, p = 0.24, ges = 0.017 for interaction). Post hoc comparisons show the main effect is mainly driven by the amblyopic group (t_17_ = -2.62, p = 0.036).

#### 3.2.5 Comparing individual differences to the severity of the amblyopic VA deficit

The amblyopic deficit is typically quantified using broadband letter acuity, either as the acuity in the amblyopic eye or the difference in acuity between the eyes. We correlated this measure against the results from our tasks where participants identified letters in limiting external noise or scrambling.

In the top row of **Figure 12**, we plot the interocular threshold ratio from Experiment 2B against the interocular broadband letter acuity ratio. We find a significant correlation only for the bandpass noise condition (**Figure 12**A). Participants with a greater amblyopic deficit (according to their amblyopic eye acuity for broadband letters) also show a deficit in the amblyopic eye’s efficiency for identifying letters in bandpass noise.

**Figure 12:**
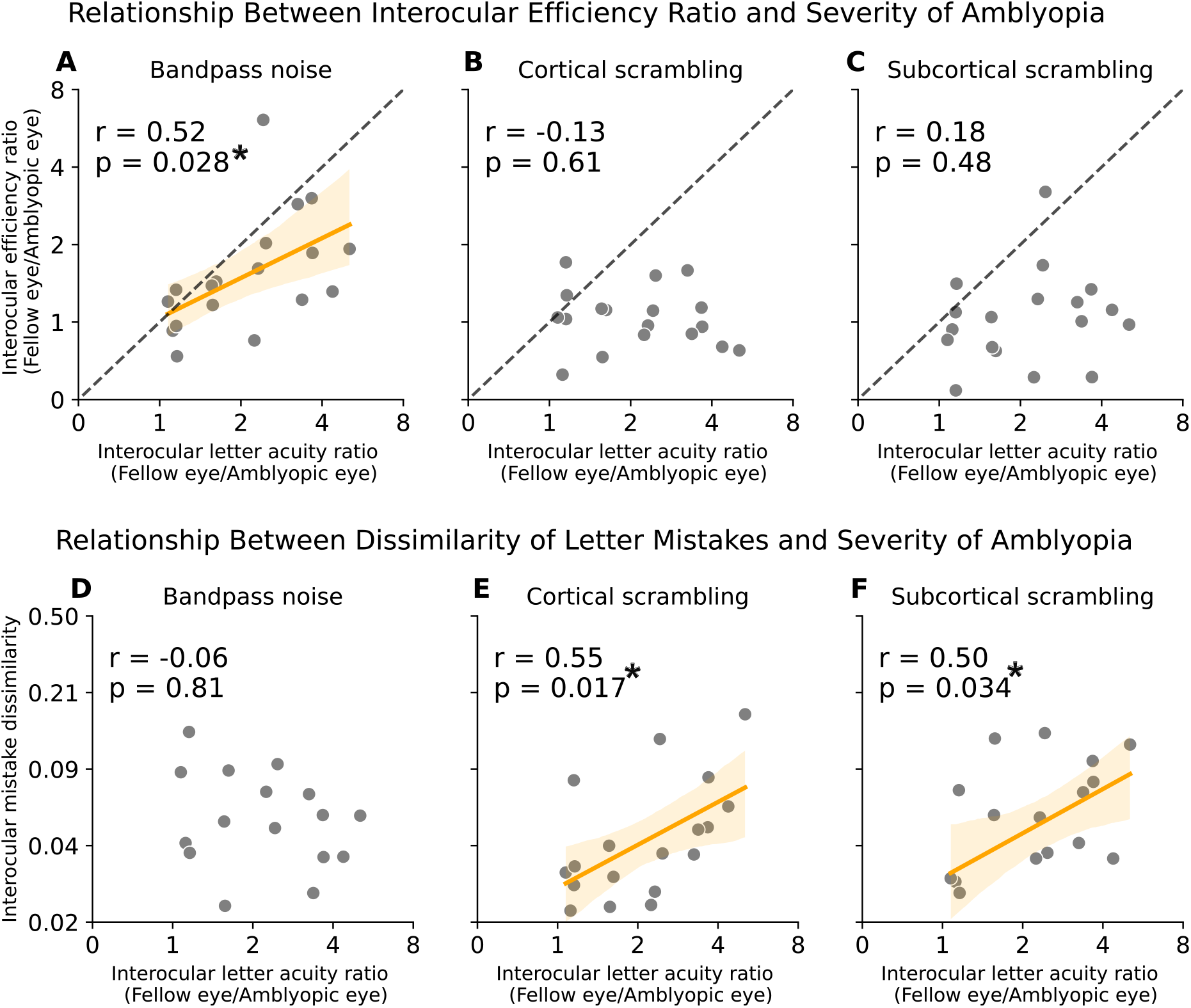
Scatter plot of interocular efficiency ratio (top row) and interocular mistake similarity (bottom row) against the severity of amblyopia. The axes are log-log scaled. Interocular mistake dissimilarity is calculated using the weighted KL divergence between the fellow and amblyopic eyes. The higher the divergence, the more dissimilar the mistakes. Correlation coefficient and associated p value are shown for each panel. A regression line is shown with 95% confidence region for cases where the p value is smaller than 0.05.

The bottom row of **Figure 12** shows the correlation between the dissimilarity of the mistake pattern between the two eyes (calculated individually for each participant) and the severity of amblyopia. Contrary to the correlations for the performance measure described above, this analysis of the mistake patterns shows a significant correlation *only* for the two scrambling conditions (**Figure 12** E-F). Although we do not find a significant performance deficit that is related to the severity of amblyopia (**Figure 12** B-C) we nevertheless see that individuals with more severe amblyopia show a distinctive pattern of mistakes in the responses made using their amblyopic eyes.

## 4 Discussion

We investigated the behaviour of amblyopic and healthy control participants performing a task that required the identification of bandpass letters. We applied both a bandpass noise mask and a physiologically-inspired scrambling algorithm to distort the letters. This allowed us to measure the threshold magnitude of noise or scrambling at which identification was still possible. We have previously established that our three stimulus conditions place different demands on visual processing such that different strategies are required to optimise performance on each task. This was done by training separate convolutional neural networks on the different conditions and comparing their performance across the entire stimulus range (Zhu et al., 2024). In this study, we again find evidence that these tasks place different demands on observer. Where we looked at computer vision previously in Zhu et al (2024), this time we look at how the normal and amblyopic visual system handle these different stimuli.

### 4.1. Performance in limiting external noise or spatial scrambling

We find that the amblyopic eye is particularly vulnerable to the effects of the bandpass noise. This was true even after compensating for the contrast sensitivity deficit (Experiment 2A) or a deficit in preferred spatial scale (Experiment 2B). The lower threshold noise for criterion performance (62.5% in our 4AFC task) means that the amblyopic eye was less efficient at extracting the letter identity from the noisy background. This would agree with the previous finding of Pelli et al., (2004), which showed the amblyopic eye has elevated contrast threshold for identifying letters embedded in white noise across a wide range of external noise levels. Our approach is not able to examine whether there is a greater intrinsic noise or scrambling in the amblyopic eye (i.e. higher equivalent input noise) as our task did not measure the full noise masking function. We will address this question in upcoming work using different task designs that allow the intrinsic scrambling to be quantified (Baldwin et al., 2016; Christensen et al., 2019; Solomon, 2009).

At the group level, we find no significant difference between the eyes for our amblyopic participants with our cortical and subcortical scrambling manipulations. However, when partitioning our amblyopic participants by the degree of their *binocular* impairment (determined by their Randot stereoacuity), we do find that the amblyopic eye efficiency is significantly greater for participants with better binocular vision (for bandpass noise and cortical scrambling in Experiment 2A, and for all three stimulus conditions in Experiment 2B). We would predict that the degree of interocular suppression would also correlate with the our measurements, as we have previously shown a relationship between interocular suppression and stereoacuity in amblyopia (Webber et al., 2020). Previous work has shown that perceptual distortions in amblyopia are associated with binocular functions (Piano et al., 2015). We previously proposed that such distortions may be responsible for the elevated equivalent internal disparity noise in amblyopia (Alarcon Carrillo et al., 2023). This could explain the efficiency difference we see between the group with good stereo and the group with poor/no stereo. We will pursue the possible relationship with spatial scrambling in future studies using a common spatially bandpass stimulus design for both tasks.

In Experiment 1A, we find a further curious result which is significant in our control group (but only appears as a trend in the amblyopic group). There is a greater efficiency for the dominant eye identifying letters affected by our “subcortical” scrambling. Broadly speaking, our approach would consider the cortical scrambling to relate to the heterogeneity of V1 neuron projections forming subfields of V2 receptive fields (Tao et al., 2014). On the other hand, the subcortical scrambling targets heterogeneity one stage below, from the monocular inputs (projecting from the lateral geniculate nucleus) to the cortex. The finding that efficiency in subcortical scrambling depends on the viewing eye in normal vision suggests that our subcortical scrambling noise may indeed target a mechanism prior to binocular combination in V1.

### 4.2. Double-pass consistency and analysis of mistake patterns

Comparison of two repetitions of the Experiment 2B where participants responded to identical stimuli (**Appendix A4**) reveals that the mistakes were partially but not entirely determined by the stimulus image the participant viewed on each trial. If the participant’s visual system were a simple linear classifier, unaffected by any noise or uncertainty of its own, then we would expect consistent responses to the same stimuli. We saw a significant, but moderate stimulus-driven consistency, that varied depending on the participant group and stimulus condition.

We analysed the pattern of mistakes made by the different participant groups using their stronger or weaker eyes. We find that participants using an amblyopic eye made a distinct pattern of mistakes, which would be consistent with spatial distortion (or possibly an under-sampling) affecting the decoding of letter identity from that eye’s input. This echoes previous behavioural findings from spatial tasks where amblyopic participants exhibit poor pooling of local stimulus features (Chandna et al., 2001; Hess et al., 1999; Kozma & Kiorpes, 2003; Mussap & Levi, 2000; Simmers & Bex, 2004).

### 4.3. Relation to the visual acuity deficit

The results of our acuity measurements show that the broadband and bandpass versions of the letter acuity task show a stronger correlation (*R*^2^ = 94%) than between either letter acuity measure and the grating acuity task (*R*^2^ = 76 to 78%). This discrepancy is part of our study’s motivation, as a significant scrambling in amblyopia could account for additional difficulty in a letter acuity task.

When relating our results from Experiments 2A back against the severity of the participants’ amblyopia (determined by their interocular VA ratio), we found a curious divergence between the results from our bandpass noise stimulus and from our two types of scrambling. For bandpass noise, there was a significant deficit in the amblyopic eye’s efficiency for performing the task. This was not present in either type of scrambling noise, but we did find the pattern of mistakes to be *more different* between the two eyes in this stimulus condition for individuals with more severe amblyopia. The association of letter mistake divergence and acuity provides new insights into the neural basis of acuity loss in amblyopia. As this result occurs only in the scrambling conditions, it suggests that internal topographical disarray could play a central role in limiting acuity.

### 4.3. Conclusion

We implemented a new stimulus design where letters were distorted by an algorithm to mimic possible physiological scrambling in the amblyopic visual system. Using this method, we showed that the tolerance for such scrambling was overall similar in the amblyopic and normal visual system when deficits in contrast sensitivity or spatial scale were accounted for. Despite the scaling by visibility or acuity, the amblyopic eye was still less efficient in bandpass noise. Furthermore, we found promising individual differences that showed significant relationships between task performance and the stereoacuity/visual acuity deficits in our amblyopia group. This included both analyses of the noise thresholds for identifying scrambled letters, and an examination of the pattern of mistakes that were made.

## Additional Information

## Acknowledgements

This work was supported by an FRQ fellowship (RXZ) and CIHR Vanier award (RXZ) to RXZ, funding from the Canadian Institutes of Health Research awarded to RFH (#8139) and applies new methods developed under a Natural Sciences and Engineering Research Council of Canada (NSERC) Discovery Grant awarded to ASB (RGPIN-2022-04216).

## Author contributions in CREDIT format

**RXZ:** Conceptualization, Data Curation, Formal Analysis, Investigation, Methodology, Software, Visualization, Writing - Original Draft. **RFH:** Conceptualization, Funding Acquisition, Supervision, Writing - Review & Editing. **ASB:** Conceptualization, Funding Acquisition, Formal Analysis, Methodology, Software, Visualization, Supervision, Writing - Review & Editing.

## Appendix

### A1. Clinical information for amblyopic participants

**Table A1:**
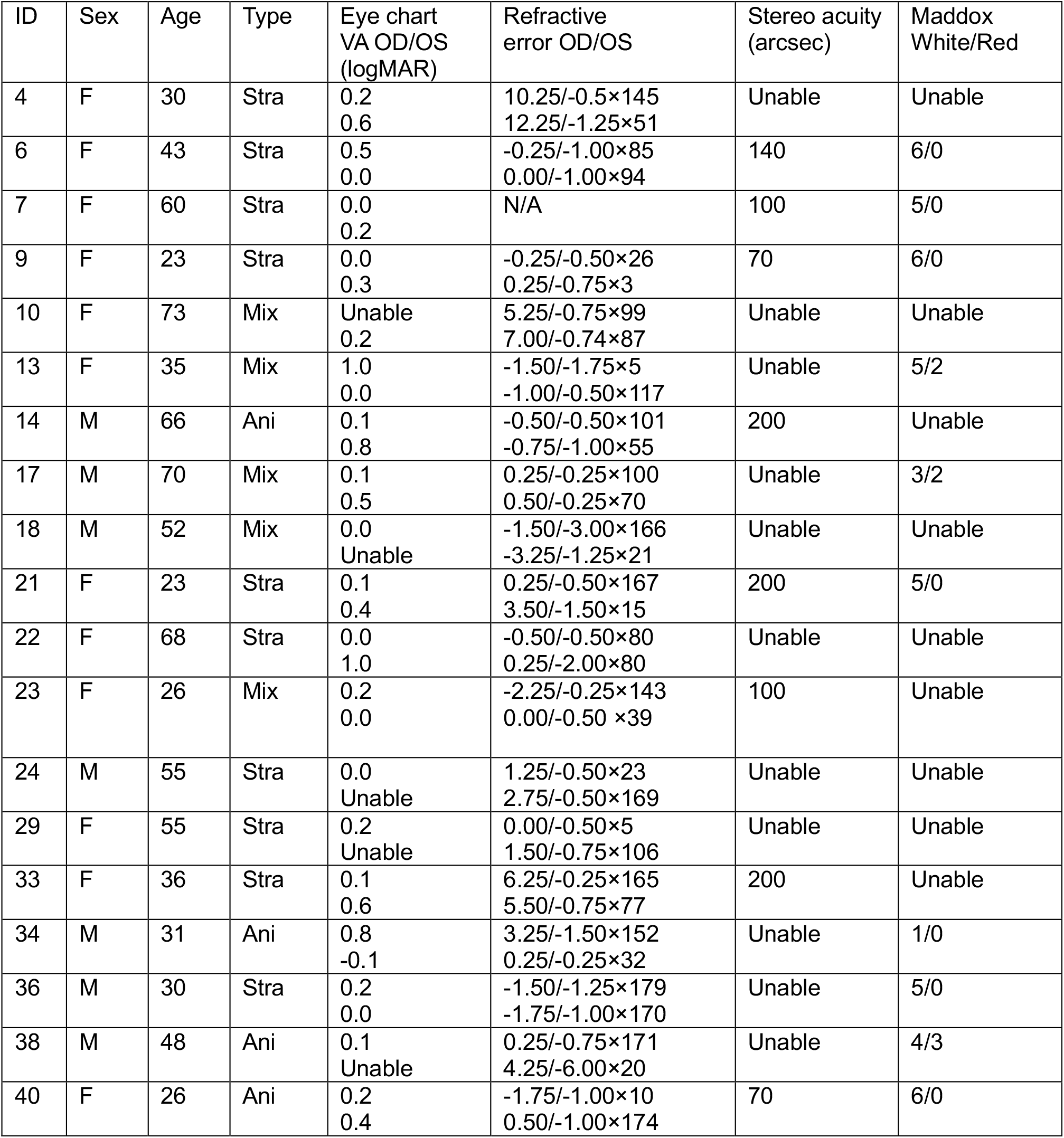
Clinical details of amblyopic participants. “Aniso” denotes anisometropia, “Mix” denotes strabismus + anisometropia, “Strab” denotes strabismus.

### A2. Confusion matrix example

**Figure A1:**
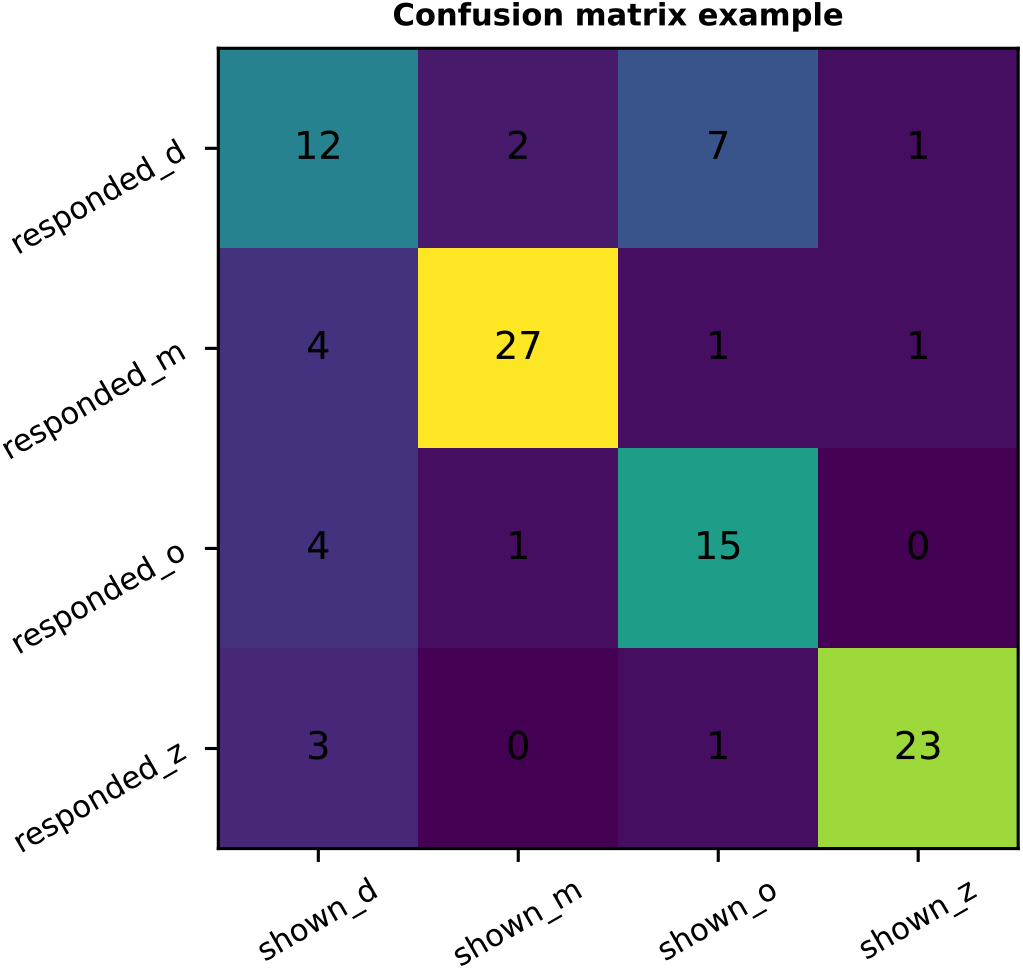
An example confusion matrix. Entries are counts. Each column represents the counts of different letter responses for each shown letter identity.

### A3. Linking deviance with KL divergence

For two vectors *p* and *q* for which we assume *p* is the probability vector measured from the data and *q* is the probability vector of the model predictions. Since the saturated model predictions are the same as *p*. The deviance of *p* from *q* becomes:

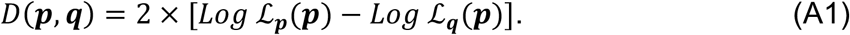

Expanding the log likelihood, we get:

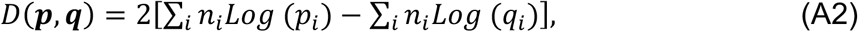

Where *n*_*i*_ is the number of times the *i* th outcomes occurs in our data and *N = ∑*_*i*_ *n*_*i*_ is the total number of trials. Simplifying the equation above further we have:

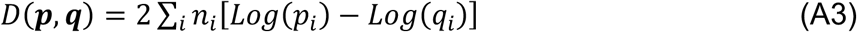

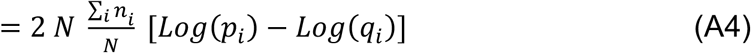

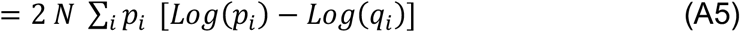

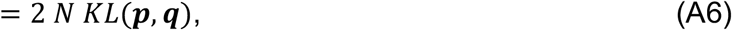

where *KL(**p, q**)* is the KL divergence.

### A4. Double-pass consistency analysis

The analysis we used to quantify the double-pass consistency is shown in **Figure A2**. In this dataset, the exact same set of stimulus conditions was shown to each participant twice. This allows us to calculate both the proportion of correct responses for each stimulus level, and the proportion of responses where the same letter identity was given as a response to the stimulus (including mistakes) in both repetitions.

We quantify consistency by comparing against a simple model where the mistake responses are random (i.e. they are not determined by the stimulus). In that case,

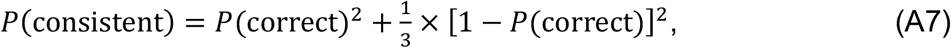

a consistent response occurs either when the response to a stimulus was correct in both repetitions, or when both responses were incorrect but the same mistake was made in both repetitions (a 1 in 3 chance). This model curve is shown in **Figure A2**A, with example data from the stronger and weaker eyes of a single participant.

To quantify the overall consistency, we calculate the area between the data points and the model curve by first transforming the data such that the model curve is a vertical line at x=0 (**Figure A2**B). Areas to the *left* of the model curve (where participants were *less* consistent than they should have been even by chance) count as negative area, meaning that the expected total area for a participant who had a random pattern of mistakes would be zero. For the maximum case where the responses were totally consistent, the area would be 0.375.

**Figure A2:**
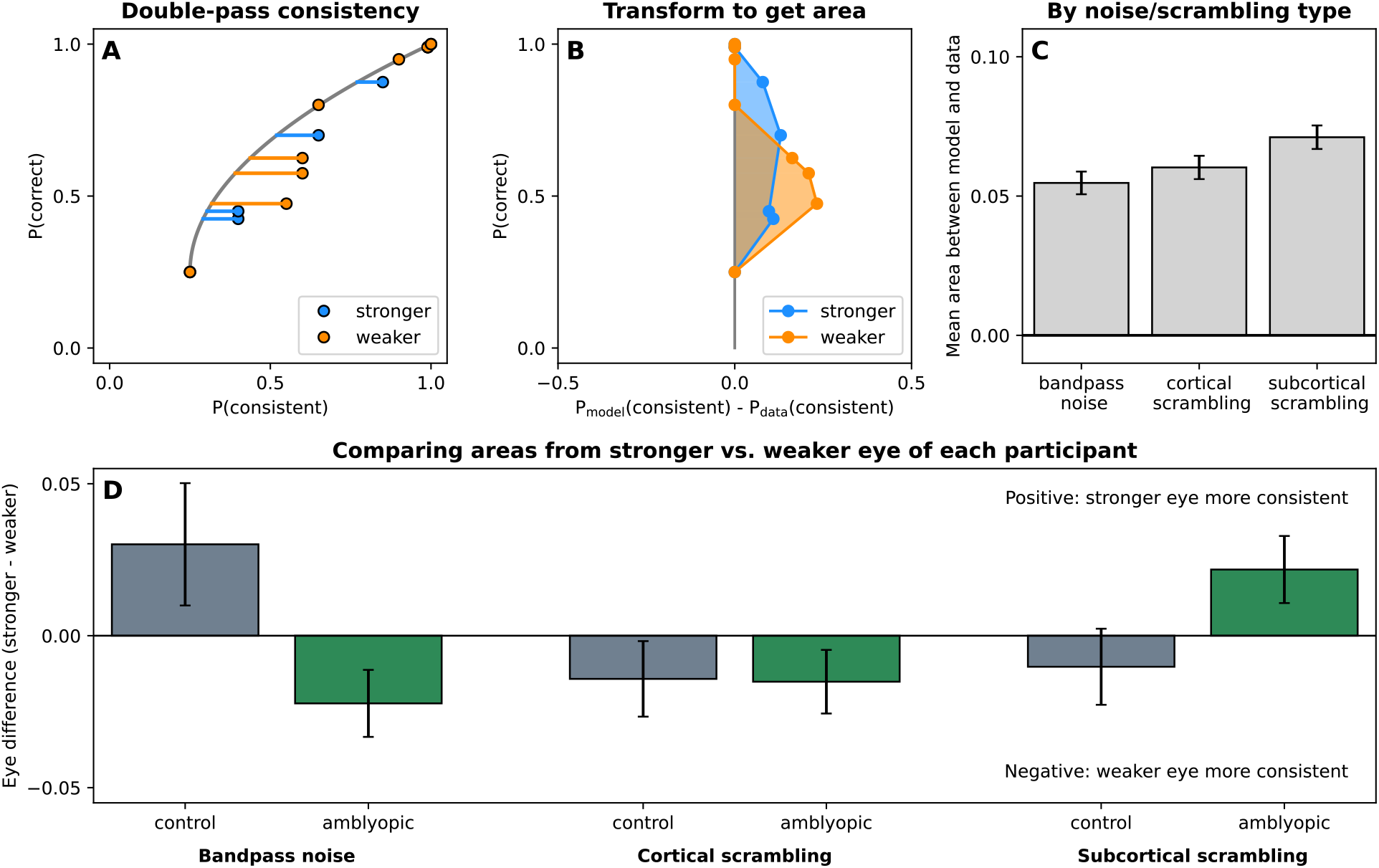
Panel A shows data from a single participant, plotting the proportion of correct responses against the proportion of responses where a consistent letter response was given. The grey model curve gives the predicted relationship for random mistakes responses (Equation A7). To quantify the overall consistency, we transform these points by subtracting the x-coordinate of the model curve (Panel B) and calculate the area between the data and the model. Panel C shows the mean area for each stimulus condition (collapsed across eyes and participant groups). Panel D compares the difference between stronger and weaker eyes for each group with each noise or scrambling condition. All error bars show standard error.

The average area for each stimulus condition (our three noise or scrambling types) is significantly above zero (**Figure A2**C), indicating that our stimuli drive letter responses that are more consistent than expected by chance. We conducted a three-way ANOVA on factors of stimulus condition, participant group (amblyopic vs control) and eye (stronger vs weaker). One outlier value was Winsorised. Comparison between the different stimulus conditions was not significant (F_2,54_ = 2.26, P = 0.114, ges = 0.02). The bottom panel (**Figure A2**D) visualises the significant three-way interaction (F_2,56_ = 5.25, P = 0.008, ges = 0.04) which is driven by a significant difference between the two participant groups in bandpass noise (t_17.39_ = -2.19, P = 0.042), bolstered by a non-significant difference in the opposite direction with the subcortical scrambling stimulus (t_24.22_ = 1.77, P = 0.090).

Looking at each participant group individually, a two-way ANOVA (factors of stimulus condition and eye) revealed a significant interaction for the amblyopic group (F_2,34_ = 3.98, P = 0.028, ges = 0.14), and a non-significant interaction for the control group (F_2,22_ = 2.87, P = 0.078, ges = 0.12). Overall, we conclude that the different stimulus conditions we compare lead to different levels of consistency in double-pass analysis when comparing the stronger and weaker eyes of our amblyopic and control participants.

